# Functional border-associated macrophages limit Alzheimer’s Disease progression

**DOI:** 10.64898/2026.01.31.703045

**Authors:** Drew Adler, Natália Pinheiro-Rosa, Alon Millet, Zoran Z Gajic, Evan Cheng, Begoña Gamallo-Lana, Yosuke Morizawa, Eric Klann, Wen-Biao Gan, Adam C Mar, Jose H. Ledo, Hernandez Moura Silva, Juan J. Lafaille

**Affiliations:** Vilcek Institute of Graduate Biomedical Sciences, New York University Grossman School of Medicine; New York, NY, USA; Department of Cell Biology; New York University Grossman School of Medicine, New York, NY, USA; Department of Neuroscience and Physiology, New York University Grossman School of Medicine; New York, NY, USA; Laboratory of Systems Cancer Biology, The Rockefeller University; New York, NY, USA; Tri-Institutional Program in Computational Biology and Medicine, The Rockefeller University; New York, NY; New York Genome Center; New York, NY, USA; New York University Grossman School of Medicine; New York, NY, USA; Department of Neuroscience and Physiology, Neuroscience Institute, New York University Grossman School of Medicine; New York, NY, USA; Center for Neural Science, New York University, New York, NY, USA; Institute of Neurological and Psychiatric Disorders, Shenzhen Bay Laboratory, Shenzhen, 518132, China; Department of Neuroscience, Department of Pathology and Laboratory of Medicine, South Carolina Alzheimer’s Disease Research Center, Medical University of South Carolina; Charleston, SC, USA; Laboratory of Immunophysiology, Ragon Institute of Mass General, MIT, and Harvard; Cambridge, MA, USA; Department of Biology, Massachusetts Institute of Technology; Cambridge, MA, USA; Howard Hughes Medical Institute; Cambridge, MA, USA; Department of Pathology, New York University Grossman School of Medicine; New York, NY, USA

## Abstract

Brain-resident macrophages are known to play numerous roles in the progression of Alzheimer’s Disease (AD). However, the relative contribution of microglia and border-associated macrophages (BAM) to AD pathogenesis has been difficult to disentangle. We recently identified *Maf*, a newly described AD GWAS gene, as essential for BAM, but not microglial, survival. By crossing BAM depleted mice with the 5xFAD AD model, we found stark evidence of cerebral amyloid angiopathy (CAA), increased overall β-amyloid burden, accelerated markers of neurodegeneration, and early memory deficits. In the healthy brain, BAM take up more β-amyloid per cell than microglia. However, as disease progresses, both in human AD patient samples and model AD mice, BAM number is reduced, and the remaining BAMs display impaired endocytic capacity, and show signs of metabolic exhaustion at an earlier age than microglia. Thus, strategies to preserve or restore BAM function represents a novel therapeutic avenue for AD and CAA.

## Introduction

Alzheimer’s disease (AD), the most common cause of dementia, is pathologically defined by the presence of extracellular deposition of amyloid-β (Aβ) and intracellular neurofibrillary tangles of hyperphosphorylated tau protein ^1,2^. Beyond clear morphological changes found in microglia (MG) from AD patient samples, myeloid cells have arisen as strong contributors to AD risk, as many genes discovered in AD gene wide association studies (GWAS) are either related to myeloid cell function or occur in myeloid cell-specific genes (>50% of total GWAS loci) ^3–9^. While phagocytosis and waste clearance is a core function of macrophages, the role of brain macrophages in Aβ clearance remains controversial, with some groups reporting an amyloid clearance role ^10–12^, others reporting a pro-amyloidogenic seeding role ^13–16^, and still others reporting a seeding role for parenchymal plaque and a clearance role for vascular plaque (the defining feature of cerebral amyloid angiopathy, CAA) ^17,18^.

Likely contributing to this uncertainty, brain resident macrophages have largely been treated as one group and referred to as microglia. While meningeal and perivascular brain macrophages have long been identified, (e.g. ^19–25^) it had been difficult to distinguish them from blood-monocyte-derived macrophages infiltrating the brain borders. Only recently have border associated macrophages (BAMs) been recognized as the resident macrophages of the meninges, perivascular spaces, and choroid plexus, as a complement to the microglia, the brain parenchyma’s resident macrophage ^26–37^. Besides their CNS border locations, BAM have been subdivided by their phenotype into BAM1 (CD206^LO^ MHC classII^HI^) and BAM2 (CD206^HI^ MHC class II^LO^)^30,32,38^.

A significant problem in the field is that the tools used to study microglia are not specific to the cell type. For instance CSF1R inhibitory drugs ^39^, genetic manipulation (*Cd11b* ^40^, *Cx3cr1* ^41,42^, *Csf1r* ^43–45^), also significantly target BAMs. Likewise, tools used to specifically deplete BAMs, such as or toxic liposome delivery ^35,46^, have resulted in a significant or total targeting of other myeloid cells ^47–50^ and other methods such as anti CSF1R antibodies rely on an intact blood-brain barrier (BBB) ^28^ which is not present in many neuroinflammatory contexts ^51,52^.

We recently discovered that the transcription factor c-MAF (encoded by *Maf*) specifically regulates transcriptional programs of vasculature-associated macrophages (VAMs) in multiple organs ^32^. In peripheral organs of mice with conditional knockout of *Maf*, VAMs remain present but are phenotypically distinct. However, in the brain, BAM2 cells are eliminated from birth and do not repopulate, without altering microglia survival ^32^ or function (see below), consistent with the finding that microglia are dependent on a related long-*Maf* family gene, *Mafb* ^53^. Importantly, *MAF* has independently arisen as one of 31 new AD GWAS genes with a higher likelihood of being a causal risk gene for AD and related dementias (out of ∼100 total AD GWAS loci identified to date) ^54^. However, the specific contribution of BAM populations to AD pathogenesis and a mechanism of MAF’s involvement in AD are presently unknown.

Herein, we leveraged BAM2 depleted mice crossed with the 5xFAD AD mouse model along with postmortem human AD samples and an *in silico* metabolic flux analysis of human BAMs to provide a temporal model for BAM involvement in AD pathogenesis. We identified c-MAF-dependent BAM2 as exceptionally effective endocytic cells, with a high capacity for Aβ uptake. We show that BAM2 are important mitigators of AD progression early in disease; however, as disease progresses, BAM2 mitochondrial function becomes impaired and BAM2 cells reduce in number, limiting their benefit. The reversion of BAM2 decline is therefore a novel path to delay the onset of AD-related cognitive impairments.

## Results

### BAM2 cells are the most phagocytic brain cells and are enriched in AD risk genes

To gain an understanding of core pathways and functions of BAMs vs microglia that might be relevant for AD, we conducted single cell-RNA sequencing (Sc-RNAseq) on subdural CD45^+^ brain immune cells isolated from wild-type (WT) adult mice (5-6 months). While other transcriptional atlases of brain immune cells have been published ^30,35,38^, many did not capture significant BAM populations, which only represent ∼4-6% of the CD45^+^ brain immune landscape. To further select for BAMs, we utilized a brain macrophage enrichment protocol (see *methods*) and filled our flow cell with all sorted CD45^Hi/Int^ cells, which are enriched in BAMs, followed by CD45^Lo^ cells, which are predominantly microglia ^32^ (**Fig. S1A,B,C**). Through this method we captured 3.6K BAMs (clusters 3-4, 14.3% of total cells) and ∼21.2K microglia (clusters 0-2, 5-6) out of a pool of 25137 cells. We defined BAM cluster 3 as BAM1, expressing high levels of MHC ClassII, *Ccr2*, and *Cd74*, and BAM cluster 4 as BAM2, expressing high levels of *Mrc1*=CD206, *Lyve1*, *Cd209f*, and *Cd163*, as we and others have previously described ^30,32,38^ (**Fig. S1A,B and Table S1**). We could further delineate BAM1 as CD45^Int^/CDllB^+^/CD64^+^/MHCII^+^, BAM2 as CD45^Int^/CDllB^+^/CD64^+^/MHCII^-^/CD206^Hi^, and microglia as CD45^Lo^/CDllB^+^/CD64^+^/MHC II^-^/CD206^-^ cells as using flow cytometry (**Fig. S1A)**

KEGG pathway analysis comparing cluster 4 (BAM2, **Fig. S1B**) to an aggregate microglia cluster (0-2, 5-6, **Fig. S1B**), revealed enrichments in the pathways “Phagosome” and “Endocytosis” (**Fig. 1A,B).** Within the “Phagosome” pathway 38 vs 11 genes were significantly upregulated in BAM2 and within “Endocytosis” 38 vs 14 genes were upregulated in BAM2 compared to microglia (**Table S1**). Considering a large number of AD risk genes are specifically related to phagocytosis ^6^, likely important for the clearance of substances that accumulate in the disease, we assessed whether BAM clusters at baseline were enriched in AD risk alleles associated with known genes compiled in the NHGRI-EBI GWAS catalogue for the trait “Alzheimer’s Disease” (MONDO_0004975). Indeed, BAM2 were more enriched in identified risk alleles than microglia (100 vs 73, **Fig. 1C and Table S1**), with particularly high levels of AD high risk genes *Apoe*, *App*, Ms4a family members, and the aforementioned gene *Maf* (**Fig. 1C**). To statistically assess whether BAM2 were truly enriched for AD risk, we generated an AD enrichment score based on the relative expression of NHGRI-EBI GWAS alleles and compared BAM to microglia aggregate clusters. Indeed, BAM2 had higher AD enrichment scores than microglia (**Fig. 1D**). Similarly, BAM2 had higher expression of endocytosis pathway genes than BAM1 (**Fig. S1D,E**), expressed more AD risk alleles (32 vs 16, **Fig. S1F and Table S1**) and had higher AD enrichment scores than BAM1 (**Fig. S1G**). Thus, gene expression data suggests that BAM2 represent a cluster of brain macrophages have a high capacity for phagocytosis and are enriched in AD risk genes.

**Figure 1.**
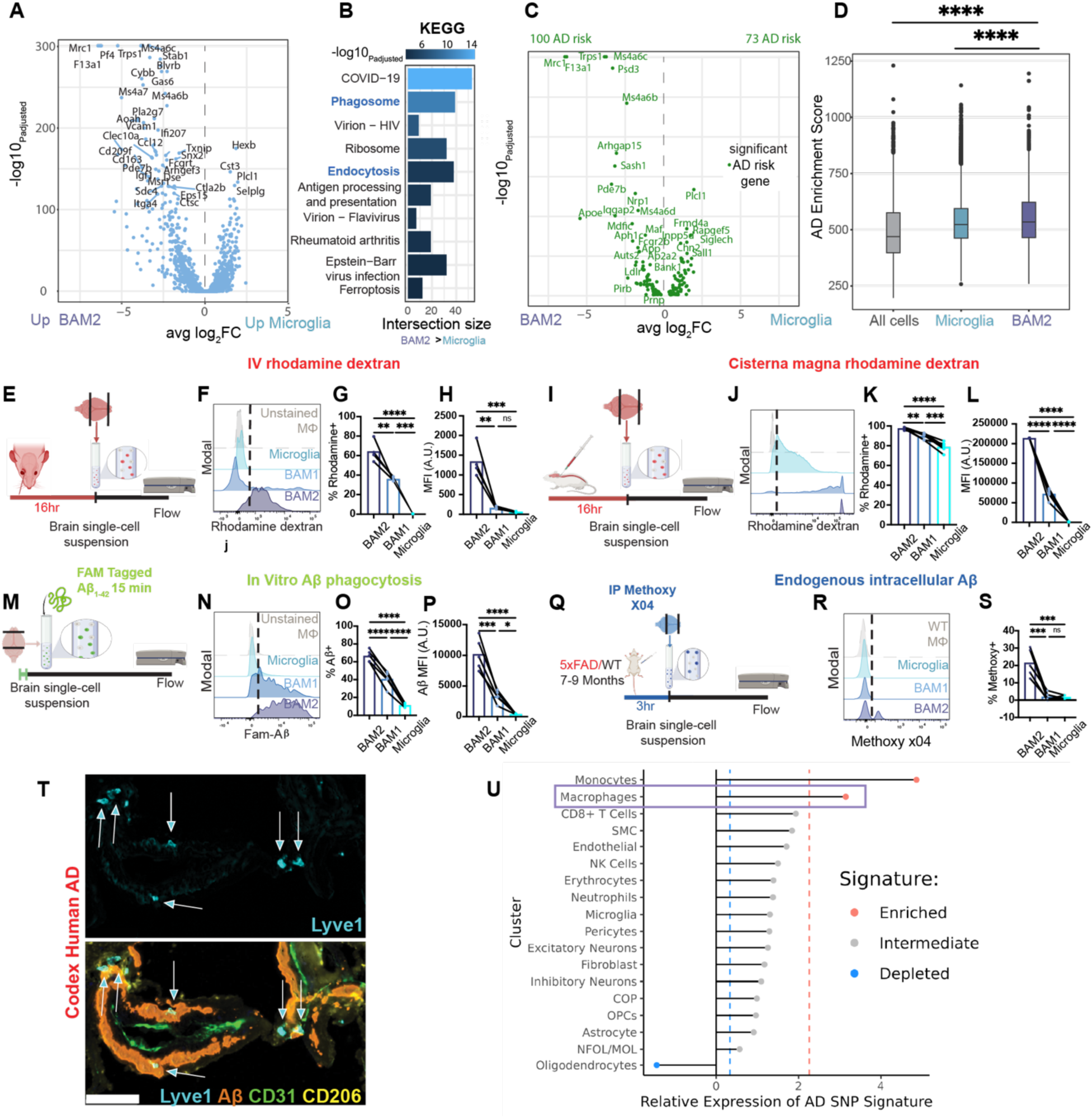
BAM2 are highly phagocytic and enriched in AD risk. **A,** Volcano plot with all significant differentially expressed genes between BAM2 (Cluster 4 in **Fig. S1B**) and microglia (Clusters 0,1,2,5,6) (n=4 WT mice 5-6 Mo, 2M, 2F). **B**, All KEGG pathways enriched in BAM2 > microglia. **C**, Similar to (**A**) but only for all mapped genes associated with an AD risk allele (MONDO_0004975). **D**, Whisker plots surrounding whisker plots with median, 25% and 75% depicting the distribution of AD enrichment score per cell within total clusters, microglia, or BAM. **E**, Experimental design for rhodamine dextran i.v. administration; **F**, representative histograms for each brain macrophage subtype; **G**, quantification of % of rhodamine positive cells; **H**, quantification of median fluorescence intensity (MFI) for intravenous rhodamine endocytosis assay (n=4 WT mice). **I-L** As with (**E-H**) but for an experiment in which rhodamine dextran was injected into CSF via cisterna magna (n=6 WT mice). **M-P,** As with (**E-H**) but for an experiment in which single cell suspensions were incubated with fluorescein amidite (FAM) tagged Aβ_1-42_ oligomers for 15 minutes (n=5 WT mice). **Q-S,** As with (**E-G)** but for an experiment in which Methoxy-X04 was injected intraperitoneally (IP) 3 hours prior to the generation of single cell suspensions (n=5 5xFAD mice + WT mouse for control comparison). **T,** Representative image with white arrows depicting LYVE1^+^CD206**^+^** BAM2 in human post-mortem superior frontal gyrus colocalized with Aβ deposited around CD31^+^ vasculature. Scale bar=50 µm. **U**, Analysis of the change in expression of human AD GWAS genes comparing AD cells to healthy cells within the same cluster (genes from MONDO_0004975, see *Materials and methods* section for datasets used). A Hampel filter was used to calculate threshold values and colored dots represent clusters enriched or depleted in change from healthy to AD. (**D**) Wilcoxon rank sum test with continuity correction. (**G-H, K-L, O-P, S**) One-way matched ANOVA with Sidak’s multiple comparisons test. *P<.05, **P<.01, ***P<.001, ****P<.0001. Bar plots with connecting lines represent populations from the same animal. AD= Alzheimer’s Disease.

To directly determine the endocytic capacity of the brain macrophage subtypes, we injected WT mice intravenously (IV) with rhodamine-dextran 70kD and assessed cellular uptake 16 hours later via flow cytometry (**Fig. 1E**). BAM2 took up far more blood-borne product than BAM1 or microglia, both as percentage of rhodamine^+^ cells and as mean fluorescence intensity (MFI) (**Fig. 1F-H**). However, the perivascular location of BAM may provide first access to blood-borne dextran. To control for this possibility and effects of the blood-brain barrier (BBB), we injected rhodamine dextran into the cerebrospinal fluid (CSF) through the cisterna magna (**Fig. 1I**). Here again, the percentage of BAM2 uptake of rhodamine-dextran was significantly higher than the other brain macrophages (**Fig. 1J-L**).

To determine whether BAM2 had an intrinsically enhanced endocytic capacity for the most synaptotoxic form of Aβ, soluble Aβ_1-42_ oligomers ^55^, we generated single cell suspensions of brain immune cells and fed fluorescently tagged Aβ_1-42_ oligomers for 15 minutes, a time measuring only phagocytic capacity and not differences in degradation ^56,57^, followed by flow cytometry (**Fig. 1M**). Again, in these primary cultures BAM2 endocytosed far more product than BAM1 (2.8x MFI) or microglia (23.5x MFI) (**Fig. 1N-P**). Finally, to determine which cell type harbors intracellularly, *in vivo*, the most fibrillary amyloid during AD, we injected intraperitoneally 5xFAD mice with Methoxy-X04, a BBB penetrant fluorescent probe that binds Aβ sheets, and assessed endogenous Aβ uptake after 3 hours ^58^ with flow cytometry (**Fig. 1Q**). Matching our *ex vivo* data, BAM2 showed a distinct second positive peak, which was absent in other brain macrophages, and significantly greater methoxy positivity (**Fig. 1R,S**). To determine whether these animal data were relevant to human disease, we stained post-mortem sections from superior frontal cortex of human AD patients. We observed a distinct overlap of LYVE1^+^ CD206^Hi^ perivascular BAMs and Aβ (**Fig. 1T**). These results identify CD206^Hi^ MHCII^Lo^ BAMs (BAM2) as remarkable endocytic cells, regardless of route of access and composition of the substance tested.

We further evaluated whether human BAMs were also notable in their differential expression of AD GWAS risk alleles between health and disease. Applying an AD enrichment score on the transcriptional delta between healthy and AD for all cell clusters identified in a recent analysis of human AD ^59^ revealed that BAMs were one of two cell clusters (of 18) with significant enrichment of AD genes, while microglia were not significantly enriched relative to the background of all other cell types (**Fig. 1U**). Together these results suggest that BAM2 carry significant known genetic AD risk, particularly in pathways related to endocytosis.

### BAM2 functional and numerical decline in Alzheimer’s disease

Continual exposure to aggregated proteins has recently been documented to induce “exhausted” or “senescent” microglial states ^57,60,61^, as high levels of phagocytosis eventually lead to exhaustion ^62^. Given the superior endocytic activity of BAM2, we hypothesized that BAM exhaustion may develop as disease progresses. To evaluate endocytic exhaustion relevant to AD, we exposed brain macrophages to fluorescent Aβ oligomers as in **Fig. 1M**, comparing WT and 5xFAD mice at a young age (3-4 months) and at a later age (7-9 months). In 3-4-month-old mice, both BAM2 and microglia *ex vivo* endocytosis of Aβ were unchanged between WT and 5xFAD mice, while maintaining the striking advantage of BAM over microglia described above (**Fig. 2A,B**). However, at 7-9 months of age, BAM and microglia diverged in their endocytic progression: significantly fewer 5xFAD BAM2 cells were Aβ-positive than WT controls, while in the same mice, slightly more 5xFAD microglia were Aβ-positive than WT controls (**Fig. 2C**).

**Figure 2.**
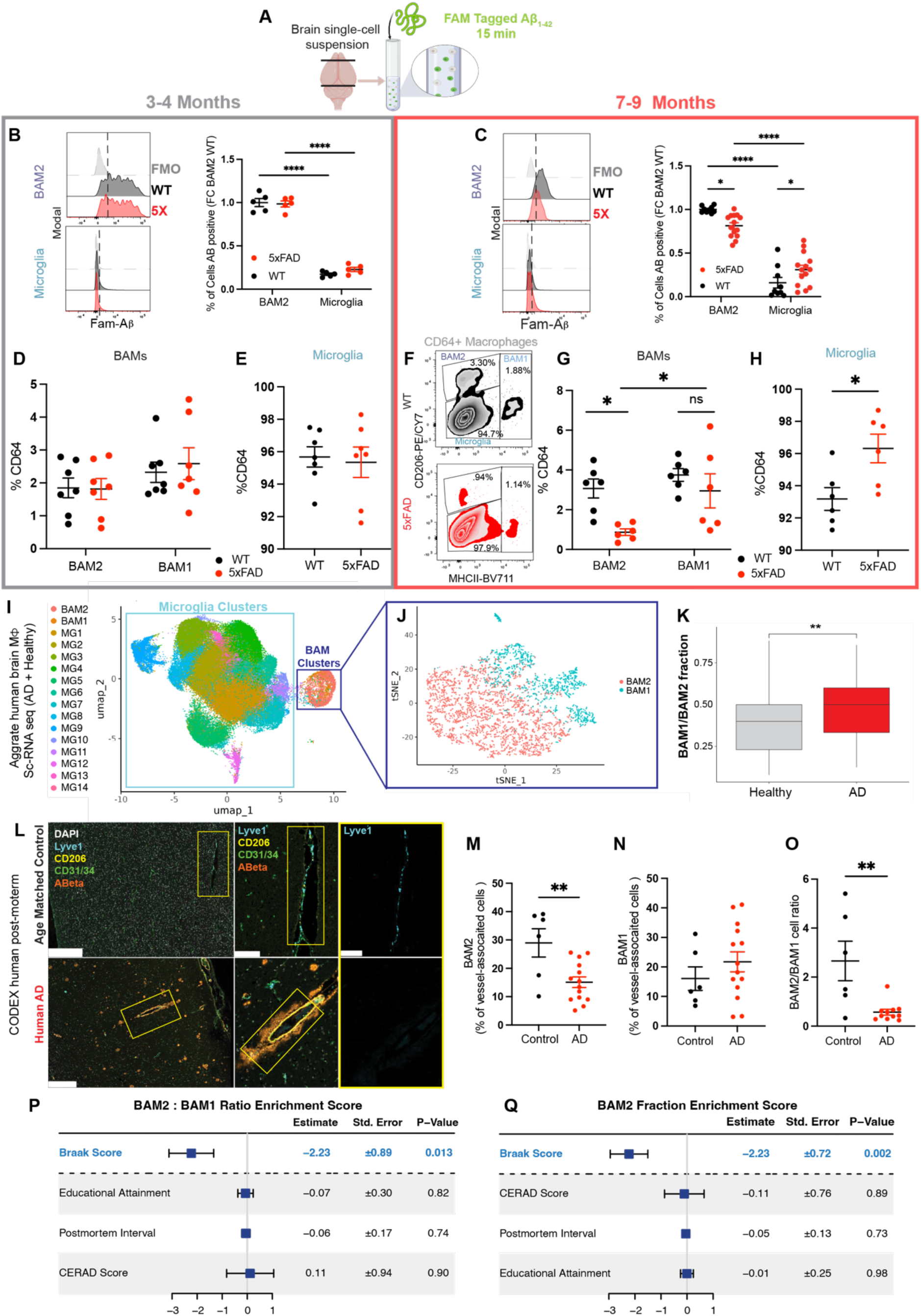
AD progression reduces BAM2 function and number. **A,** Schematic of oligomeric soluble Aβ_1-42_ endocytosis assay (as in Fig. 1M); **B**, representative histograms of Aβ_1-42_ positivity (right of dotted line) for BAM2 (top) and microglia (bottom) with respective quantification of fold change in 3-4 mo mice (n= 5 WT, 5 5xFAD mice); **c**, as with (**B**) but for 7-9 mo mice. WT avg (n=9 WT, 14 5xFAD mice). **D-E**, Quantification of (**D)** BAM and (**E**) microglia as percentage of CD64^+^ (total) macrophages (n=6 WT, 7 5xFAD mice); (**F)** representative zebra plots from 7-9 mo mice from WT (top, black) and 5xFAD (bottom, red) mice with percentage of each brain macrophage subtype; **G-H,** as with (**D-E**) but for 7-9 mo mice (n=8 WT, 6 5xFAD mice); **I**, UMAP of aggregated human brain macrophages from healthy control and AD patients highlighting microglia clusters (light blue box) and BAM clusters (dark blue box). **J**, UMAP of BAM subclusters. **K**, BAM1/BAM2 fraction per patient from the ROSMAP dataset (n=151 healthy, 81 AD patients). **L**, Representative micrographs of CODEX-stained AD and age-matched control superior frontal gyrus for BAM2 maker LYVE1, CD206, vasculature maker CD31 and CD34, and Aβ along with large vessel ROIs. **M-O**, quantification of the (**M**) % of vessel associated cells that are BAM2 (LYVE1^+^), (n=6 healthy, 14 AD), (**N**) BAM1 (CD206^+^LYVE1^-^), (n=6 healthy, 14 AD) and (**O**) the ratio of BAM2/BAM1 (LYVE1^+^/CD206^+^LYVE1^-^) (n=6 healthy, 11 AD). **P,Q**, Forrest plots demonstrating correlation between (**P**) BAM2/BAM1 enrichment score and different AD-clinical measures. (**Q**) % of BAM2s of total macrophages and different AD-clinical measures. (**B,C,D,G**) Two-way matched ANOVA with Sidak’s multiple comparisons test. (**E,H,K,M,N,O**) Two-tailed unpaired t-test. *P<.05, **P<01, ****P<.0001. (**i**) Scale bars=500µm (left), 100 µm (middle/right). Fold change normalized to 1 (as average) includes individual values that are not 1; these derive from littermates with the same genotype and slightly different individual values. UMAP= uniform manifold approximation and projection.

Next, we asked if disease progression only reduced the functionality of BAM2 cells or whether there was also a decline in cell number. As with microglia, BAM2 are long-lived, yolk-sac-derived macrophages without significant replacement from bone marrow derived monocytes under regular circumstances ^34,38^. We determined that at 3-4 months of age there was no difference between WT and 5xFAD mice in the proportion of any of the three brain macrophage populations out of the pool of CD64^+^ macrophages (**Fig. 2D,E**). However, at age 7-9-months, the proportion of BAM2 cells among CD64^+^ macrophages decreased in 5xFAD mice, while the proportion of microglia increased in the same mice (**Fig. 2F,G,H**). Thus, the microglia:BAM2 and BAM1:BAM2 ratios increase as disease progresses.

To determine if these changes in mouse BAM2 number extended to human AD, we generated a molecular atlas of brain macrophages that included ∼4000 human BAMs from healthy and AD patients from datasets comprising cells from 513 patients derived from the Religious Orders Study and Rush Memory and Aging Project (ROSMAP) ^63,64^ (**Fig. 2I**). Subclustering BAMs by excluding microglia clusters revealed two distinct clusters of BAMs (**Fig. 2J**), generating a human BAM subtype signature (**Table S2**). As with our GWAS analysis on human brain cells, we noticed that AD profoundly changed the transcriptional landscape of human BAMs to a greater extent than microglia (**Fig. S2**). We then determined whether the BAM clusters significantly changed in proportion between control and AD brains. Although we didn’t identify a disease-associated BAM cluster as has been described for microglia ^64^, we found that the MHC class II^+^ cluster (BAM1) was significantly enriched in brains from AD patients, while the BAM2 cluster was reduced (**Fig. 2K**

To corroborate these findings *in situ*, we stained amyloid rich superior frontal gyrus from human post-mortem AD patients and age matched controls for BAM2 marker LYVE1, CD206, endothelial markers CD31 and CD34, and Aβ using CODEX ^65^ multiplexed immunofluorescence. We trained a neural network using HALO AI to identify penetrating arterioles and post capillary venules, where perivascular BAMs reside ^34^, and automatically quantified the number of LYVE1^+^ (CD206^Hi^, BAM2) and CD206^+^LYVE1^-^ (MHC II^+^, BAM1) as a percent of blood vessel-associated cells. Here, as with our results in 5xFAD mice, we found a strong decrease in BAM2 cells, unchanged BAM1, and a concomitant decrease in the BAM2/BAM1 ratio (**Fig. 2L-O**). These results suggest that amyloidogenic conditions are associated with reduced BAM2 function and number, which may contribute to disease progression.

Finally, to see if BAM2/BAM1 ratio or BAM2 fraction (of total macrophages) correlated with any clinical AD covariates, we merged ROSMAP clinical data with our annotated single cell dataset to generate a unified structure with both cell annotation and patient of origin metadata and then built linear models in which either BAM2:BAM1 enrichment score or BAM2 fraction are predicted by a linear combination of select clinical variables. Using this approach, we found both BAM2/BAM1 ratio and BAM2 fraction were anti-correlated with Braak score, a canonical measure of neurofibrillary tangle pathology from 0-6 (each additional Braak score is predicted to decrease BAM2 fraction by >2 fold), (**Fig. 2P,Q**). These results suggest that amyloidogenic conditions are associated with reduced BAM2 function and number and that AD pathological severity correlates with reduced BAM2 proportion.

### BAM2 depletion accelerates AD and causes marked cerebral amyloid angiopathy (CAA)

Considering the enrichment of AD risk alleles held in BAM2 and their high capacity for Aβ uptake, we hypothesized that BAM2 depletion would alter key pathological and behavioral hallmarks of disease. To explore this hypothesis, we crossed BAM2 depletion mice ^32^ (Csf1r^Cre/+^ *Maf*^F/F^) with *Maf*^F/F^ 5xFAD mice, to generate BAM depletion AD model mice (Csf1r^Cre/+^ *Maf*^F/F^ 5xFAD, hereafter Csf1r^Cre/+^5xFAD) mice. Conditional deletion of *Maf* dramatically reduced the number of BAM2 without affecting the number of microglia (**Fig. 3A**). The increased percentage of BAM1 cells does not indicate a conversion from BAM2 to BAM1. At newborn stages, both BAM2 and BAM1 are greatly reduced in Csf1r^Cre/+^*Maf*^F/F^ mice. However, unlike BAM2 which are not reconstituted from monocytes, BAM1 are reconstituted over time, showing a rebound ^32^. As further validation, we immunostained for BAM2 markers CD206 and LYVE1 and pan macrophage marker IBA1 in adipo-clear cleared brains in Csf1r^Cre/+^*Maf*^F/F^ mice and *Maf*^F/F^ littermates, and quantified BAM and microglia numbers in 4K microns of cortical tissue. *Maf* deletion led to a ∼80% reduction in LYVE1^+^CD206^+^ BAM2 without reducing the number of IBA1^+^ microglia (**Fig. 3B-F**). We also tested the functionality of microglia in BAM depleted mice. Unlike what is seen in TREM2 and TAM receptor (*Mertk* and *Axl*) knockout mice ^10,66^, microglia from Csf1r^Cre/+^5xFAD mice had unchanged endocytosis capacity for Aβ_1-42_ oligomers (**Fig. 3G-I),** as well as unchanged association with Congo Red derivative X-34^+^ plaques **(Fig. S3A-C).**

**Figure 3.**
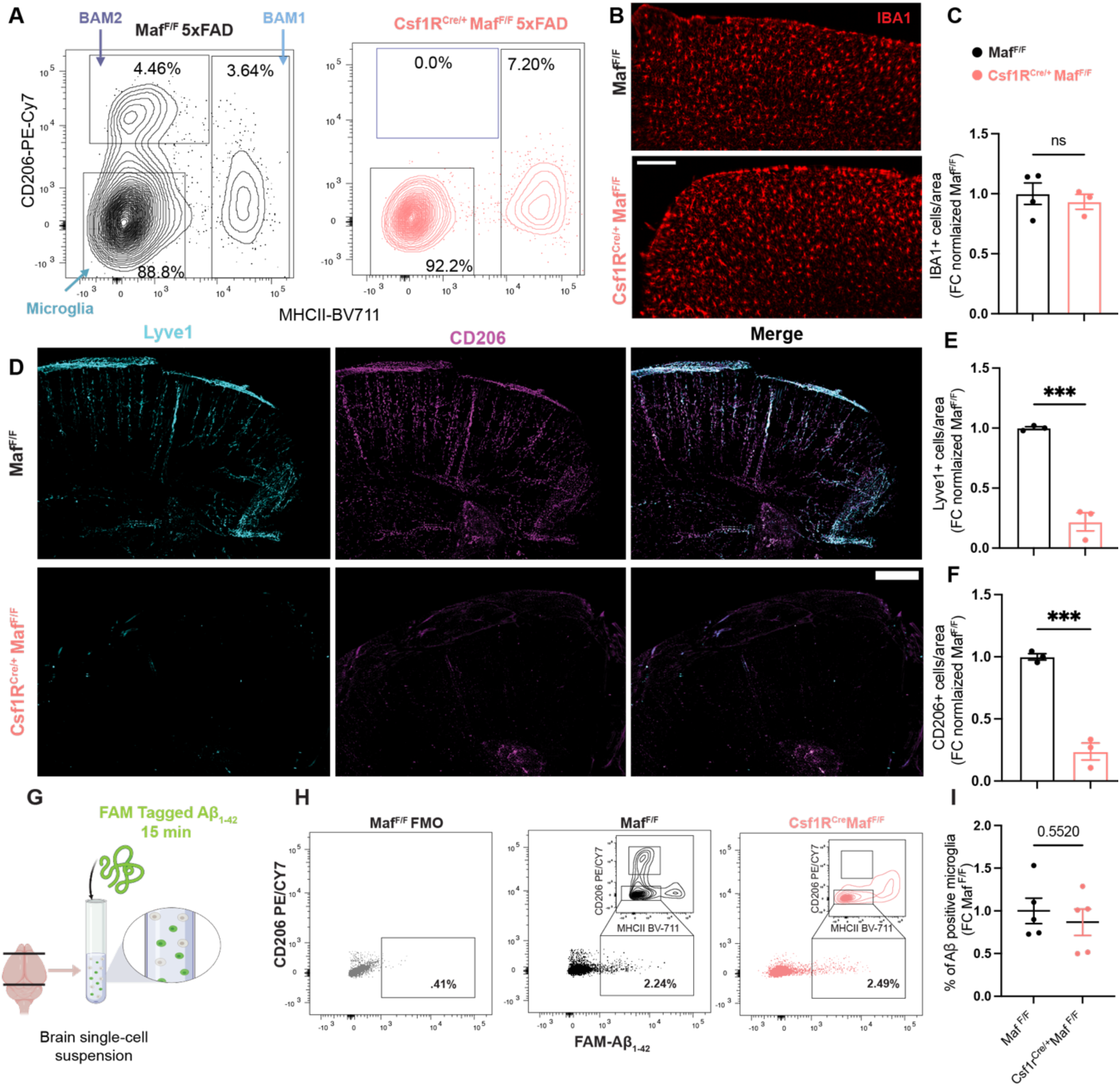
Generation of BAM2 depleted AD mice. **A,** Representative contour plots of CD64^+^ cells and relative proportions of microglia (bottom left), BAM2 (top left), and BAM1 (right) in Cre^-^ (BAM2 normal, black) and Cre^+^ (BAM2 depleted, orange) mice. **B**, IBA1^+^ macrophages (predominantly microglia, but also BAM) from adipo-cleared cortical brain tissue from BAM depleted and control mice with **C**, respective quantification. **D**, as with (**B**) but staining LYVE1^+^ and CD206^+^ BAM. **E,F,** quantification of (**E**) LYVE^+^, and (**F**) CD206^+^ BAM/area normalized to Cre^-^ average value (n=3 Cre^-^, 3 Cre^+^ for **E,F** and n=4 Cre^-^ for **C**). **G**, schematic of experiment (as in Fig. 1M) in which single cell suspensions were incubated with FAM-labeled Aβ_1-42_ for 15 minutes prior to flow cytometry. **H**, representative dot plots depicting microglia population (inset bottom square) FAM positivity between control and BAM2 depleted mice (inset top square shows absence of BAM2 in depleted mice) with (**I**) respective quantification (n=5 mice per group). Bar plots=mean +/- S.E.M. (**C,E,F,I**). ***P<.001 Two-tailed unpaired t-test. (**B,D**). Scale bars=200µm. Fold change normalized to 1 (as average) includes individual values that are not 1; these derive from littermates with the same genotype and slightly different individual values.

We next assessed whether the absence of BAM2 accelerated amyloid related pathology in the 5xFAD mouse model. We chose two age ranges to assess histopathological differences between BAM depleted mice and littermates: 3-5 months, when amyloid first starts to spread in the 5xFAD model, prior to overt differences and thus representing “pre-clinical AD”, and 7-9 months of age, after a significant buildup of plaque and clear memory deficits representing overt “clinical AD.” ^67,68^. To control for established age, sex, and recently identified inheritance-based differences in the model ^69^, we expressed all differences as fold change over same sex littermates. At 3-5 months Csf1r^Cre/+^5xFAD mice had significantly greater deposition of fibrillary amyloid (as indicated by X-34 staining ^70^) in both cerebral cortex (3.0 fold) and hippocampus (6.7 fold) (**Fig. 4A,B,C**). Even starker, while at 3-5 months 5xFAD mice with normal BAMs had virtually no CAA (0.6% of total large vessel area), in Csf1r^Cre/+^5xFAD mice, nearly 5% of blood vessel surface was covered in plaque (>7 fold over littermates) (**Fig. 4A,D**). When we carried out our analysis on 7-9 month old mice, no longer there was a difference between cortical, hippocampal, or vascular buildup of fibrillary amyloid between Csf1r^Cre/+^5xFAD and their littermates, suggesting that the above-mentioned numerical and functional decay of BAM2 (**Figure 2**) reduced their impact on disease outcome at more advanced ages (**Fig. 4A,B,C,D, and Fig. S4A**). The data expressed as fold change reflects the expected age-related disease progression in 5xFAD controls which, by 7-9 mo, catch up to BAM2-depleted mice regarding amyloid pathology.

**Figure 4.**
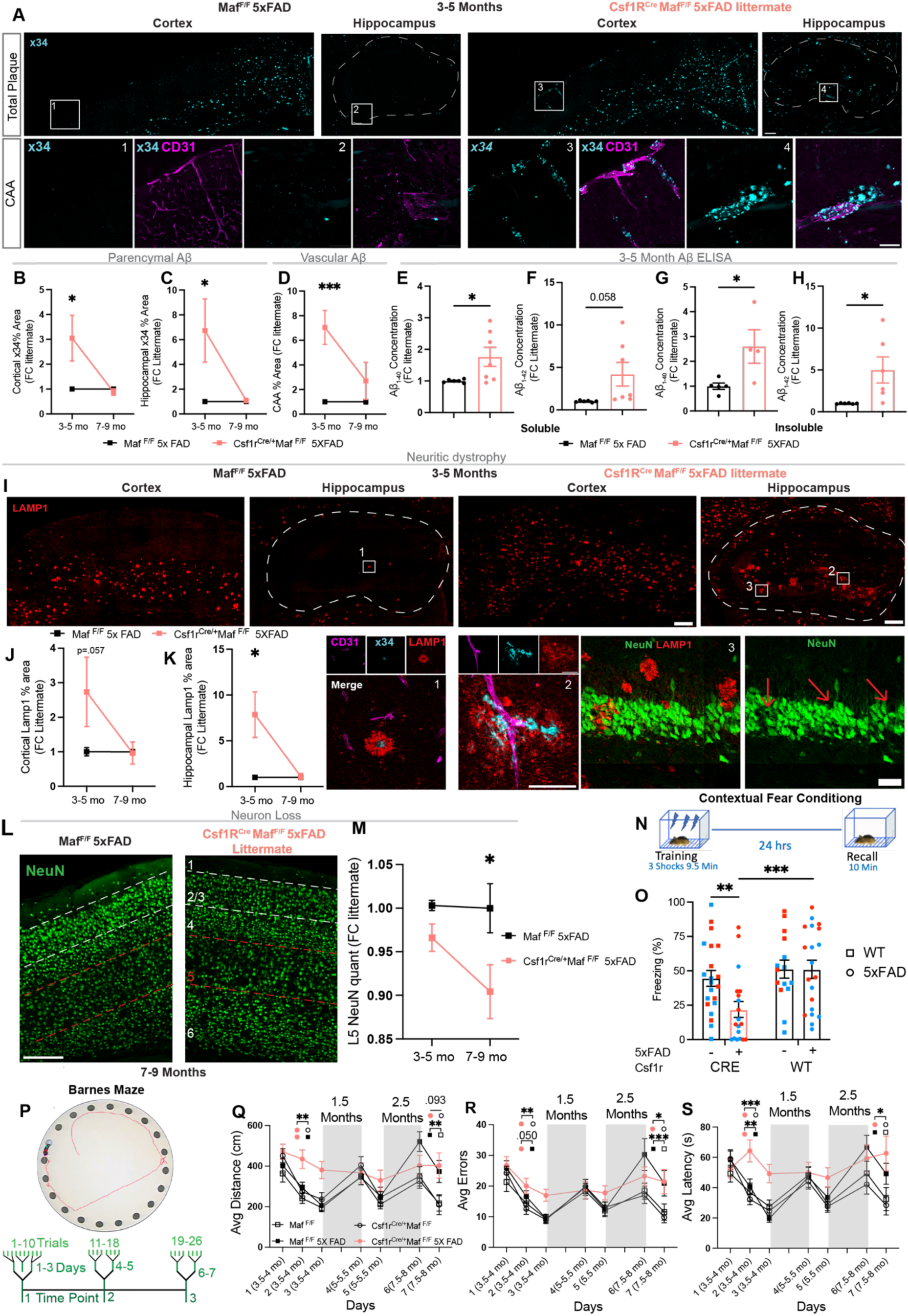
BAM2 depletion accelerates AD-related phenotypes. **A**, Representative micrographs of X-34 amyloid staining in cortex and hippocampus in 3-5 month-old Cre^+^ 5xFAD (BAM2 depleted) and Cre^-^ (BAM2 sufficient) 5xFAD littermates (top) with regions of interest (ROIs, numbered boxes below) depicting pial and penetrating CD31^+^ blood vessels in cortex and the hippocampal artery in hippocampus. **B**, Quantification of X-34 cortical area fraction normalized to littermates at 3-5 mo (n= 11 Cre^-^, 12 Cre^+^ mice) and 7-9 mo (n=7 Cre^-^, 6 Cre^+^ mice). **C**, as with (**B**) but for hippocampus. **D**, Quantification of the percentage of pial and penetrating blood vessel area covered with CAA, presented as fold change normalized to littermates at 3-5 mo (n= 6 Cre^-^, 6 Cre^+^ mice) and 7-9mo (n=7 Cre^-^, 6 Cre^+^ mice) mice. **E-H**, ELISA based quantification of 3 mo whole brain (**E)** soluble Aβ_1-40_ (n= 6 Cre^-^, 7 Cre^+^ mice), (**F)** soluble Aβ_1-42_ (n= 6 Cre^-^, 7 Cre^+^ mice), (**G**) insoluble Aβ_1-40_ (n= 5 Cre^-^, 4 Cre^+^ mice), and (**H**) insoluble Aβ_1-42_ (n= 6 Cre^-^, 6 Cre^+^ mice). **I**, representative micrographs of LAMP1 staining in cortex and hippocampus in 3-5 mo Cre^+^ and Cre^-^ littermates, along with high magnification ROIs (numbered boxes below) in sections. Insets 1 and 2 show sections from Cre^+^ mice depicting LAMP1 localization around X-34^+^ plaque (Inset 1) and vasculature (Inset 2, showing salient CAA). Inset 3 depicts CA3 granular cell layer in hippocampus from Cre^+^ mice in which LAMP1 staining appears in gaps in NeuN+ neuron staining (red arrows). **J**, Quantification of cortical LAMP1 area fraction at 3-5mo (n=10 Cre^-^, 9 Cre^+^ mice) and 7-9mo (n=7 Cre^-^, 7 Cre^+^ mice); **K**, as with (**J**) but for hippocampal LAMP1 at 3-5mo (n=9 Cre^-^, n=12 Cre^+^ mice), and 7-9mo (n= 6 Cre^-^, 5 Cre^+^ mice). **L-M**, neuronal loss in Cre^+^ mice. (**L**) representative micrograph depicting NeuN+ neuron staining in cortical layers from 9 mo Cre^+^ and Cre^-^ littermates; **M**, quantification of L5 neuron number/area normalized to littermates at 3-5 mo (n=6 Cre^-^, 6 Cre^+^ mice) and 7-9 mo (n=8 Cre^-^, 6 Cre^+^). **N-O,** behavioral tests. **(N),** schematic of contextual fear conditioning paradigm; (**O)**, quantification of freezing proportion in first 5 minutes of memory retrieval (blue=male (M), red=female (F), n=8 WT_M_, 7WT_F_, 11 Cre^+^_M_, 10 Cre^+^_F_, 11 5xFAD_M_, 9 5xFAD_F_, 9 Cre^+^ 5xFAD_M_, 10 Cre^+^ 5xFAD_F_). Main effect of sex (F(1,67)= 10.133, p = 0.002) and Cre status (F(1,67)= 10.012, p = 0.002). **P**, Schematic of Barnes maze test along with longitudinal design including trials, days, and age (months). **Q-S** Quantification, Cre x 5xFAD x Age interaction, and 5xFAD main effect respectively of (**Q**) average distance traveled (F(6,1855) = 2.978, p = 0.007) and (F(1,1855) = 6.200, p = 0.013); (**R**) committed (F(6,1856) = 3.267, p = 0.003) and (F(1,1856) = 12.550, p < .001); and (**S**) latency (s) to locate the escape hole for each day and age (F(6,1856) = 2.098, p = 0.051) and (F(1,1856) = 8.447, p = 0.004)). See *Materials and Methods* for n values). (**A, I**) White dotted line represents hippocampal area. (**B-D, J-K, M, O**) Two-way ANOVA with Sidak’s multiple comparisons test, (**E-H**) two-tailed unpaired t-test, (**Q-S**) Linear Mixed Effects Model with Sidak’s multiple comparisons test. Significant pairwise comparisons within time points indicated by shapes associated with respective genotypes in the panel legend. *P<.05,**P<.01, ***P<.001, ****p<.0001. (**A,I,L**) Scale bars=200µm large images, 50µm high mag ROIs. Fold change normalized to 1 (as average) includes individual values that are not 1; these derive from littermates with the same genotype and slightly different individual values.

We further tested whether the buildup of Aβ seen in young BAM2 cell-depleted mice with histology was amyloid species or solubility dependent, as soluble oligomeric forms of Aβ are thought to be more synaptotoxic ^55,71–73^, and the longer Aβ_1-42_ (vs shorter Aβ_1-40_) is more associated with AD ^74,75^. After removing olfactory bulbs and cerebellum, which do not have amyloid early in disease ^67^, we performed diethylamine and formic acid extraction ^56^ to isolate soluble and insoluble fractions of brain homogenate in 3-month-old Csf1r^Cre/+^5xFAD mice and their littermates. Quantifying soluble and insoluble Aβ_1-42_ and Aβ_1-40_ using ELISA revealed significant increases in all but one (soluble Aβ_1-42_, p=.058) of these forms of Aβ (**Fig. 4E,F,G,H**). Together these results indicate that depletion of BAM2 leads to increases in Aβ buildup early in disease, irrespective of species, solubility, or location.

The amyloid cascade hypothesis contends that the deposition of Aβ sets off a cascade of events including tau hyperphosphorylation, neurofibrillary tangles, and eventually, synapse loss and overt neurodegeneration ^76,77^. This tau phosphorylation has also been shown in the 5xFAD mouse model of AD ^78^, a primary amyloid buildup model. The connection between amyloid buildup and tau appears to be through formation of LAMP1^+^ dystrophic axon spheroids, which grow around amyloid, serve as seeding loci for tau, and block action potential propagation ^79,80^. We therefore assessed whether BAM2 depletion led to an increase in dystrophic axons by performing LAMP1 staining at the two age ranges included in our analyses. As previously reported ^80,81^, we noticed distinct LAMP1 staining surrounding fibrillary X-34^+^ plaques. Mirroring X-34 staining, there was a significant increase in LAMP1 area in Csf1r^Cre/+^5xFAD mice compared to BAM-sufficient 5xFAD littermates in younger mice, but not at later ages (**Fig. 4I,J,K, and Fig. S4B**). Of further note, we observed dramatic increases in LAMP1 staining immediately adjacent to vascular-associated Aβ in BAM2-depleted mice (**Fig. 4I**, **inset 1 and 2**) and LAMP1 staining distinguishable in apparent holes in the dense hippocampal neuron granular cell layer (NeuN^+^), suggestive of potential neurodegenerative phenomena (**Fig. 4I, inset 3)**

In the 5xFAD mouse strain, overt neuron loss does not occur until ∼9 months of age, after the beginnings of cognitive impairment, and is most apparent in cortical layer 5 (L5) and the subiculum, where intraneuronal Aβ is highest ^67,82^. Therefore, we tested whether there were differences in cortical L5 neuronal number between Csf1r^Cre/+^5xFAD mice and BAM-sufficient 5xFAD littermates at both age ranges. Contrary to amyloid and neuritic dystrophy quantification, we found a non-significant trend toward a decrease in L5 neuron number in 3-5 month Csf1r^Cre/+^5xFAD mice. However, in 7-9 month-old animals we perceived a significant 10% loss in L5 cortical neuron number (**Fig. 4L,M**). These results are consistent with a model in which constitutive BAM2 depletion in Csf1r^Cre/+^5xFAD mice accelerates Aβ deposition early in disease, leading to neuritic dystrophy, which results in the late-materializing neuronal loss.

Given a degree of remaining expression of *Maf* in microglia, we employed a second Cre driver for *Maf* depletion (LysM^Cre^) to validate these phenotypes. *Lyz2* (LysM) has ∼27 fold higher expression in BAM2 than microglia, is a documented poor microglial Cre driver ^38,83^, and strongly reduces BAM2 number in *Maf*^F/F^ mice (**Fig. S5A and S5B**). The specificity of the BAM2 depletion versus microglia is consistent with the expression of different MAF family members: *Maf* mRNA is significantly enriched in BAM2 over microglia (2.3 fold) while microglia highly express and rely on a different *Maf* family member, *Mafb* ^53^(2.5 fold higher than BAM2) **(Table S1)**. We used these LysM^Cre/+^5xFAD mice to repeat the analysis of fibrillary amyloid buildup and CAA at the 3-5 month timepoint. The results matched those obtained in the Csf1r^Cre/+^5xFAD model (**Fig. S5C-F**). Together, these results suggest that BAM2 cells are vital for curbing pathology early in disease but become less important concomitantly with their functional and numerical decline.

Finally, we asked whether the acceleration in AD histopathological hallmarks identified in BAM2-depleted mice was associated with earlier conditioned fear and spatial memory deficits - two prominent types of learning and memory impairments documented in 5xFAD mice ^84,85^. At 4.5-5 months of age, a stage before associative learning and memory deficits typically emerge in BAM-sufficient 5xFAD mice ^84^, we tested WT (*Maf*^F/F^), 5xFAD (*Maf*^F/F^5xFAD), BAM2-depleted (Csf1r^Cre/+^ *Maf*^F/F^), and BAM2-depleted 5xFAD mice (Csf1r^Cre/+^ *Maf*^F/F^ 5xFAD) in a contextual fear conditioning paradigm (**Fig. 4N**). Planned, pairwise comparisons revealed BAM2-deficient Csf1r^Cre/+^5xFAD mice froze significantly less than BAM2 sufficient 5xFAD and BAM2 depleted controls on the memory retrieval trial, while there was no difference between BAM2 sufficient 5xFAD mice and WT littermates (**Fig. 4O and Fig. S6A**). We also found significant main effects of sex and Cre status: with female mice freezing more than males, consistent with prior findings^86^, and with BAM2-sufficient freezing more than BAM2 depleted mice. These results suggest that BAM2 depletion impacts fear memory, accelerating memory decline in the 5xFAD model in the absence of BAM2s.

To further investigate the temporal progression of memory decline in BAM2-depleted mice, we conducted the Barnes Maze spatial memory test in a longitudinal design, evaluating the same mice across three ages (3.5-4 months, 5-5.5 months, and 7.5-8 months) (**Fig. 4P**). Linear mixed model analyses revealed significant Cre x 5xFAD x Age interactions on distance, errors, and a trend for latency (p=.051) to find the target escape hole (**Fig. 4Q-S and Fig. S6B-G**). As with fear conditioning, pairwise comparisons indicated that BAM2 depletion accelerated the impairments of 5xFAD mice: at 3.5-4 months old, BAM2-depleted 5xFAD mice made significantly more errors, traveled significantly longer distance, had significantly longer latency to find the escape hole, and found significantly fewer holes than BAM2-sufficient 5xFAD, or BAM2-sufficient and BAM2-depleted WT control mice (**Fig. 4Q-S and Fig. S6C-F**). Moreover, BAM2-depleted 5xFAD mice did not significantly improve their latency and ability (“found”) to find the escape hole across days, unlike all other groups. BAM2-sufficient 5xFAD mice only showed significant deficits relative to BAM2-sufficient WT controls by 7.5-8-months of age, with their performance deteriorating such that they were no longer significantly different from BAM2-depleted 5xFAD mice. As expected, 5xFAD mice showed significantly greater impairment than WT controls across all ages and outcome measures. These results indicate that BAM2 depletion accelerates both the onset and progression of learning and memory deficits in 5xFAD mice. By later stages, BAM2-sufficent mice exhibited comparable declines, consistent with our histopathological data (**Fig. 4A-K**) and the observed loss of BAM2 (**Fig. 2**) during disease progression.

### BAM2 cells become metabolically exhausted in AD mice

We next sought to find a mechanism to explain the functional and numerical changes of BAM2 cells in the context of AD which could contribute to the 5xFAD model’s accelerated decline at later timepoints. Metabolic adaptation is closely tied to brain macrophage function in the context of neurodegeneration ^57,61,87,88^, and in microglia is thought to regulate endocytosis of Aβ ^61^; however, as discussed above, BAMs are often extracted together with microglia. Considering that BAM2 endocytosis of Aβ far surpasses that of microglia *ex vivo* and *in vivo*, and that BAM2 become dysfunctional at a relatively early age, we hypothesized that these main characteristics would reflect differences in the metabolic pathway utilization and flux between microglia and BAM in health and AD.

To determine baseline mitochondrial function in the different brain macrophage subtypes, we measured mitochondrial membrane potential (ΔΨ) in 3-4 month old WT mice using charged dye tetramethylrhodamine ethel ester (TMRE) ^89^, which we detected by flow cytometry (**Fig. 5A**). At baseline, BAM2 had 35% and 52% higher TMRE MFI than microglia and BAM1 respectively, and an increased proportion of cells with high levels of TMRE than microglia, suggestive of higher mitochondrial oxidative function (**Fig. 5B,C,D**). Next, we assessed whether disease progression affected TMRE in BAM2 and microglia at two ages in which we saw either a high BAM2 function (3-4 months) or a diminished function (7-9 months). Similarly to the Aβ endocytosis assay, at 3-4 months of age we saw no TMRE differences between WT and 5xFAD mice in BAM2 cells or microglia (**Fig. 5E**). However, in 7-9 month-old mice, there were significantly fewer TMRE^Hi^ cells among BAM2, with no appreciable change in microglia TMRE (**Fig. 5F**). This suggests a disease-dependent decline in mitochondrial oxidative metabolism in BAM2 cells.

**Figure 5.**
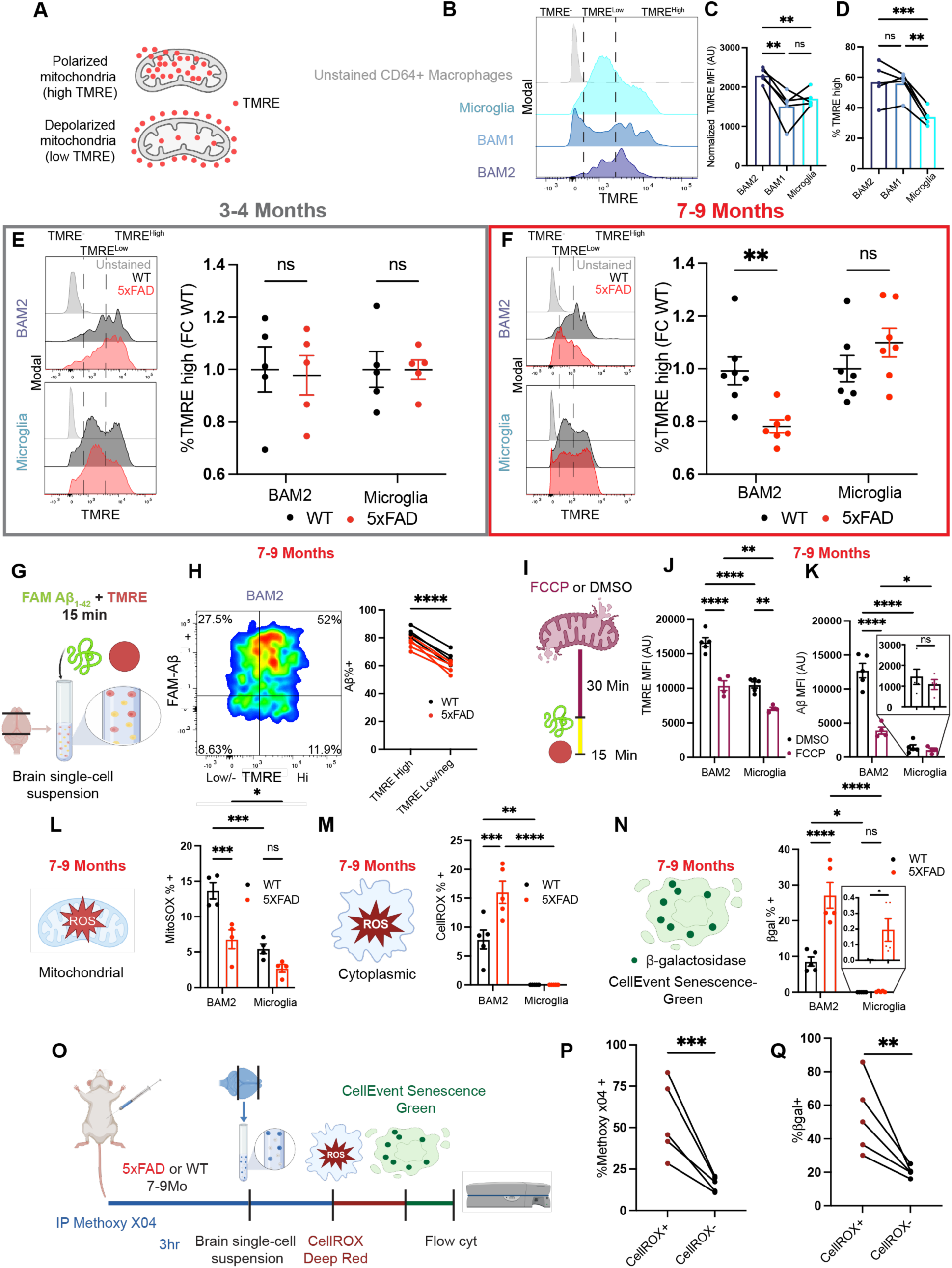
BAM2 metabolic exhaustion in AD mice. **A**, Schematic of TMRE assay. **B**, Representative histogram of TMRE staining in macrophage subtypes along with TMRE level delineation (dotted lines). **C-D**, quantification of (**c**) TMRE MFI per cell type and (**D**) percentage of TMRE^hi^. **E**, representative histograms of TMRE populations within BAM2 (top) and microglia (bottom) from 3-4 mo mice, along with respective quantification of percentage of TMRE^hi^ population normalized to avg WT levels for each cell type (n=5 WT, 5 5xFAD mice). **F,** as with (**E**) but for 7-9 mo mice (n=7 WT, 7 5xFAD mice). **G**, Schematic of experiment in which TMRE and Aβ_1-42_ were co-incubated prior to flow cytometry. **H**, representative heatmap comparing TMRE (x axis, high vs low/negative) to Aβ positivity (y axis, positive vs negative) in BAM2 cells, and respective quantification of Aβ positivity within each TMRE population (n=4 WT, 5 5xFAD mice). **I**, Schematic of experiment in which single cell suspensions were incubated with FCCP or DMSO (control) prior to Aβ_1-42_ and TMRE. **J-K**, Quantification of (**J**) mean fluorescence intensity (MFI) of TMRE and (**K**) intracellular FAM-Aβ_1-42_ for each cell type in experiment **I,** with microglia MFI magnified in inset. **L**, Quantification of percentage of BAM2 and microglia positive for mitochondrial superoxide (MitoSOX Red) from 7-9 mo mice (n=4 WT, 4 5xFAD mice). **M**, Quantification of percentage of BAM2 and microglia positive for cytoplasmic ROS using CellROX Deep Red probe in 7-9 mo mice (n=5 WT, 5 5xFAD mice). **N**, Quantification of percentage of senescent BAM2 and microglia using CellEvent^TM^ Senescence Green probe in 7-9 mo mice (n=5 WT, 5 5xFAD mice) with microglia percentage magnified in inset. **O-Q**, (**O**) Schematic of experiment multiplexing Methoxy-X04, CellROX, and CellEvent^TM^ with respective quantification (**P**) Percentage of Methoxy-X04 positive BAM2 cells correlative to CellROX positive or negative BAM2, and (**Q**) Percentage of Senescent (CellEvent^TM^ positive) BAM2 cells correlative with CellROX positive or negative BAM2 from 7-9 mo mice (n= 5 5xFAD mice). (**C,D**) One-way matched ANOVA with Sidak’s post hoc test. (**E-F, J-N**) Two-way matched ANOVA with Sidak’s multiple comparisons test. (**P-Q**) Two-tailed paired t-test. (**K,N** insets) Two-tailed unpaired t-test *P<.05, **P<.01, ***P<.001, ****P<.0001. (**C-D, H, P-Q**) connecting lines=same animal. βgal= β-galactosidase, ns= not significant, ROS/ROX= reactive oxygen species, SOX=superoxide, FCCP= carbonyl cyanide-p-trifluoromethoxyphenylhydrazone, TMRE= tetramethylrhodamine, ethyl ester. Fold change normalized to 1 (as average) includes individual values that are not 1; these derive from littermates with the same genotype and slightly different individual values.

To determine if there was an association between mitochondrial health and Aβ uptake in BAM2 cells, we measured TMRE and Aβ endocytosis simultaneously using flow cytometry, which revealed a strong correlation between TMRE level and Aβ endocytic capacity (**Fig. 5G,H**). Finally, to directly test whether active oxidative phosphorylation is required for Aβ phagocytosis in brain macrophages, we performed our Aβ phagocytosis assay in single cell suspensions preincubated with the mitochondrial uncoupling agent FCCP (**Fig. 5I**). FCCP incubation strongly reduced TMRE signal in both BAM2 cells and microglia compared to cells treated with DMSO alone (**Fig. 5J**). Intriguingly, while FCCP strongly blocked Aβ endocytosis in BAM2 cells (3.3-fold reduction in phagocytic cells), the mitochondria uncoupler had no appreciable effect on microglia Aβ endocytosis (**Fig. 5K**), suggesting that, at this stage, microglia can rely on alternative sources of ATP such as glycolysis, while BAM2 cannot. These results emphasize a dependency on mitochondrial oxidative metabolism for BAM endocytosis of Aβ and indicate basic metabolic differences between the two main brain macrophage types.

To explore a potential connection between Aβ endocytosis and cell death/exhaustion in BAM2 cells that might explain the BAM2 loss as AD progresses, we separately measured mitochondrial and cytoplasmic oxidative stress as well as cell senescence in 7-9 month 5xFAD mice using flow cytometry. Given that mitochondrial superoxide is one of the largest contributors of cellular reactive oxygen species (ROS) in macrophages ^90,91^, we measured levels of mitochondrial superoxide using MitoSOX. BAM2 cells displayed higher MitoSOX reactivity than microglia in WT mice, and BAM2 cells from 5xFAD mice had lower MitoSOX reactivity than those from WT mice (**Fig. 5L**). The production of mitochondrial superoxide is known to be primarily driven by the flux through the electron transport chain ^90–92^; thus, our results are consistent with higher baseline levels of mitochondrial membrane flux in BAM2 cells vs microglia, and a decrease in BAM2 flux observed in 5xFAD mice. Beyond mitochondria, extra-mitochondrial generation of ROS during phagocytosis via NADPH oxidase is particularly important in phagocytes where it is used in the oxidative burst to kill pathogens ^93^. Notably, of the two NADPH oxidase catalytic subunits (*Cybb* and *Cyba*), in healthy brains BAM2 cells express 13.4-fold higher *Cybb* (NOX2, P91-PHOX) than microglia (**Table S1**), while the expression of *Cyba* (P22-PHOX) is constitutive. Using the CellROX cytoplasmic ROS flow cytometry probe, we could detect cytoplasmic ROS in 7.9% of BAM2 in WT mice while microglial cytoplasmic ROS was undetectable (**Fig. 5M**). In BAM2 from 5xFAD mice, CellROX positivity more than doubled while we again could not detect cytoplasmic ROS in microglia (**Fig. 5M**) from the same mice. Finally, to determine whether BAM2 cells from 7-9 month-old 5xFAD mice showed signs of canonical senescence, we measured β-galactosidase activity by flow cytometry. We detected extremely few (<.01%) senescent microglia cells in WT mice and a modest (8.6%) number of senescent BAM2 cells (**Fig. 5N**). In 5xFAD animals the percentage of senescent BAM2 cells jumped to 27% while that of microglia slightly increased to 0.2% (**Fig. 5N**). Thus, BAM2 cells from 7-9 month-old mice at baseline produce more cytoplasmic ROS and are phenotypically more senescent than microglia, findings that are exacerbated in the context of AD.

To determine whether there was a direct *in vivo* association between the endocytosis of Aβ, oxidative stress, and senescence phenotypes that we had observed BAM2 cells from 7-9 month-old mice, we multiplexed CellROX and CellEvent Senescence probes in single cell suspensions of 5xFAD mice injected with Methoxy-X04 to label endogenous Aβ (**Fig. 5O**). We found a strong association between Aβ uptake *in vivo* and both cytoplasmic ROS and cellular senescence in BAM2 cells (**Fig. 5P,Q**). These results suggest that BAM2 cells that are actively involved in Aβ uptake produce high levels of ROS and adopt senescent phenotypes that eventually may contribute to death and dysfunction. Importantly, this is a BAM2 phenotype not shared with microglia from the same mice.

### Metabolic changes in BAM2 cells are a feature of human AD

We next examined whether features of BAM metabolic exhaustion found in the 5xFAD model extended to human AD BAMs. To assess metabolic pathway changes in human brain macrophages in AD, we employed Compass ^94^, a recently developed *in silico* metabolic flux balance analysis that can be used to accurately predict cellular metabolic states with single-cell resolution across 7,440 metabolic reactions comprised in the Recon2 consensus “metabolic reconstruction” ^95^. Feeding our human brain macrophage single-cell atlas from healthy control and AD patients into Compass generated predictive changes in metabolic flux through each Recon2 reaction along with an associated effect size (Cohen’s d statistic) for each macrophage subtype comparing AD to healthy conditions within and between cell types (**Fig. S7A and Table S3**). Each reaction was further classified into a subsystem representing a metabolic pathway with at least 3 core reactions (for instance: glycolysis/gluconeogenesis). Comparing microglia versus BAM2 from AD patients, we observed a notable divergence in the pathways enriched in each cell type.

We detected 12 and 25 pathways enriched in microglia and BAM2 respectively, out of 90 total (**Fig. S7B**). Specifically, microglia from AD individuals were enriched in sugar metabolic pathways including “Fructose/mannose metabolism,” “galactose metabolism,” and “glycolysis/gluconeogenesis”, which is consistent with our murine FCCP data that indicated that microglia, unlike BAM2 from the same brains, have the flexibility to use glycolysis when oxidative phosphorylation is absent, a capacity that BAM2 cells appear to lack. In contrast, BAMs were enriched in lipid metabolic pathways including “Cholesterol metabolism,” “fatty acid synthesis,” and “steroid metabolism” (**Fig. S7B**).

Comparing BAM2 from healthy vs AD-afflicted individuals using Compass revealed 27 (of 90) altered pathways (|Cohen’s d|>0.1 (**Fig. 6A**). This comparison revealed that flux through “Oxidative phosphorylation” and “ROS detoxification” pathways were each in the top 3 most depleted pathways in BAM2 from AD patients (versus BAM2 from healthy), matching TMRE decreases, and ROS increases found in 5xFAD mice (**Fig. 6A**). In contrast to BAM2 cells, comparing microglia from healthy vs AD patients revealed only 5 altered pathways (**Fig. 6B**). As a more statistically rigorous assessment of degree of metabolic change between cells from healthy and AD individuals, we calculated the average Euclidean distance between the computed Compass reaction penalties from cells derived from AD and healthy individuals within each macrophage subtype, which revealed that metabolic changes induced by AD in BAM2 surpass those in microglia (P_adjusted_<.001) (**Fig. 6C**). Furthermore, comparing BAM2 (B) and microglia (M) flux through individual reactions in OxPhos (**Fig. 6D**) and ROS detoxification (**Fig. 6E**) revealed predicted flux decreases from health to AD were stronger in BAM2 cells than microglia for every reaction. Thus, human BAM2 cells appear to have a greater degree of metabolic dysregulation than microglia in AD with particularly robust changes in oxidative metabolic and ROS detoxification pathways.

**Figure 6.**
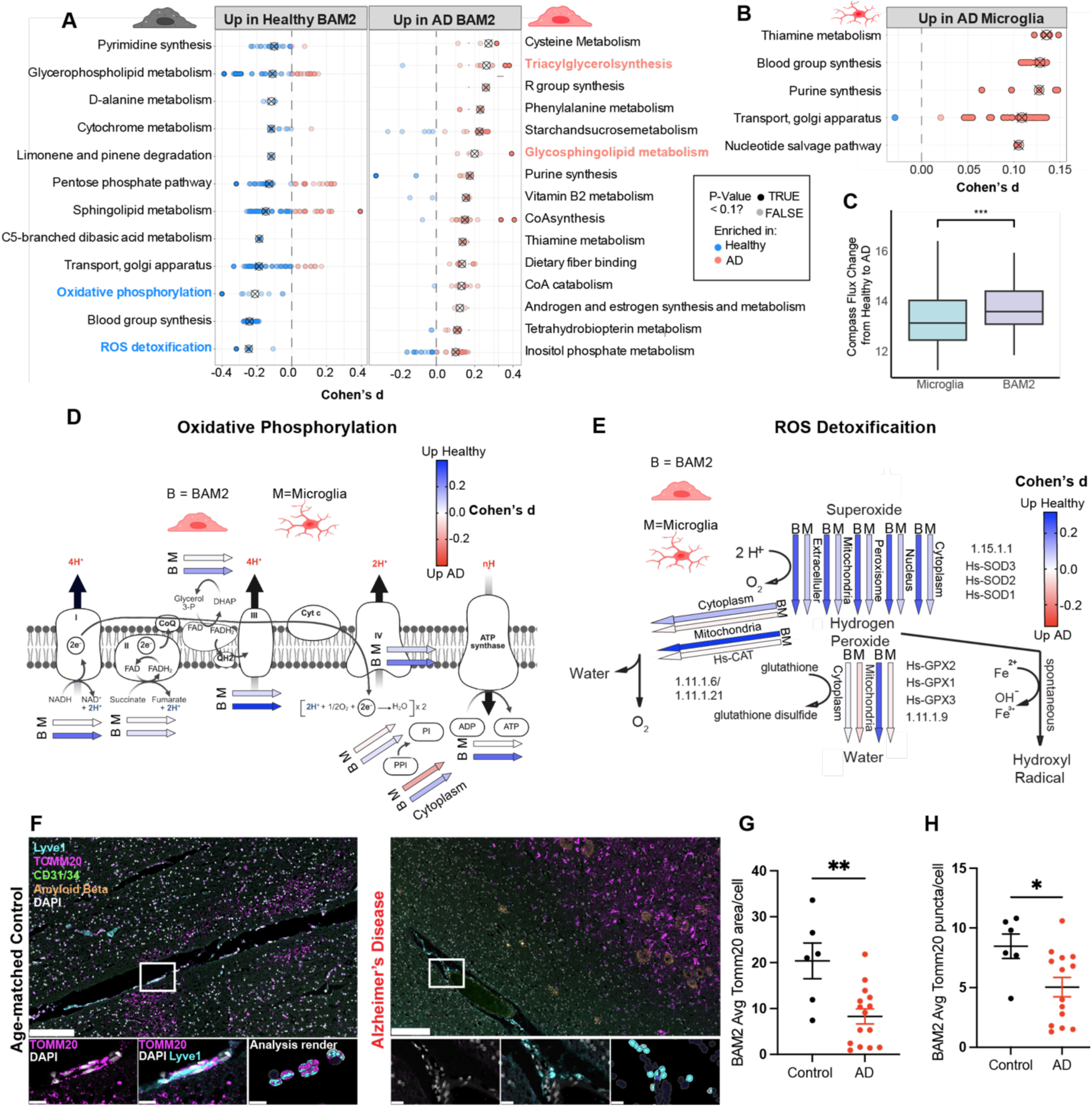
BAM2 metabolic changes in human AD. **A-B**, All COMPASS^54^ discovered subsystems (Cohen’s d>.1) up and down for (**A**) human BAM2 and (**B**) microglia comparing each cell type from AD and healthy patients (no down pathways in microglia met the inclusion criteria). **C**, Comparison of compass flux change from health to AD for microglia and CD206^Hi^ BAM (BAM2). Box plot=median value with 25 and 75 percentiles. **D-E**, All individual reactions in subsystems (**D**) “Oxidative phosphorylation” and (**E**) “ROS detoxification” with colored arrows representing Cohen’s d value in heatmap. **F-H**, (**F**) Representative images of TOMM20 mitochondrial staining in LYVE1^+^ BAMs from superior frontal gyrus of AD patients and age-matched control with quantification of (**G**) average TOM20 area per cell and (**H**) average number of TOMM20 puncta per cell (n=6 controls, 15 AD patients). (**A**) Colored subsystem names represent pathways discussed in main text. Circles with x=median enrichment for the respective subsystem, individual circles=individual reactions within the subsystem that when shaded dark =P_adjusted_<.1 (Benjamini-Hochberg adjusted). (**C**) Two-tailed paired t-test. (**G-H**) Two-tailed unpaired t-test. *P<.05, **P<.01, ***P<.001. B= BAM2s, M=Microglia.

To study mitochondrial dysregulation in human BAM2, we stained human postmortem control and AD superior frontal gyri for BAM markers and the outer mitochondrial membrane protein TOMM20 using CODEX. TOMM20 is an essential component for the mitochondrial import of cytoplasmic-synthesized proteins needed for oxidative phosphorylation, as well as mitophagy ^96,97^. Supporting our results from 5xFAD mice and Compass, we detected a strong (2.5-fold) reduction in TOMM20 reactivity as well as significantly fewer (6.5 +/- 2.0) TOMM20 puncta per BAM2 in AD samples vs age matched controls (**Fig. 6F-H**).

Together, these results support a model in which MAF-dependent BAM2 are a subset of brain macrophages that are highly metabolically active, allowing for a remarkable endocytic capacity which is highly protective against AD and CAA. However, as disease progresses, such high activity cannot be sustained, leading to declining BAMs, phenotypically and numerically, which may contribute to disease progression.

## Discussion

The advent of single-cell RNA sequencing has ushered in a new understanding of the diversity of immune cells in the brain and the way these adaptable cells change in the context to disease. Indeed, in the past decade, definitions of “disease-associated microglia,”^98^, “disease inflammatory microglia”^99^, “proliferative-region-associated microglia”^100^, “activated response microglia”^101^ “interferon responsive microglia,”^102^, “cytokine response microglia” ^103^ and “terminally inflammatory microglia”^57^ have arisen, as have two discrete border-associated macrophage subtypes, BAM1 and BAM2 ^30,32,38^. Here, by using the first genetically-based conditional BAM depletion model, we have characterized the particular importance of BAM2s in the context of AD. Having established selective BAM2 dependence (vs microglia) on the transcription factor cMAF for survival, we in turn provided a novel mechanism for the involvement of this new AD GWAS loci ^54^: *Maf* insufficiency leads to BAM2 depletion, which causes amyloid (and likely other waste product) buildup and the acceleration and increased severity of disease.

While the exact contribution of Aβ to the pathogenesis and progression of AD remains debated, it is clear that the mutations causing familial AD are directly involved in amyloid precursor protein processing into Aβ pathogenic peptides. The efficacy (albeit modest) of FDA-approved Aβ targeted immunotherapies (lencanamab, donanemab, and aducanumab) for sporadic AD unambiguously demonstrates a role for Aβ in disease ^76^. Many questions remain: for example, where does deposited Aβ have its true impact? Recent work demonstrates that the buildup of Aβ in blood vessels (CAA) is particularly nefarious as it constricts capillary bloodflow ^104^, limits perivascular waste elimination ^105^ and causes microbleeds and infarcts ^106^. In this regard, another question that our work might shed light on is the individual difference among patients between the impact on neuronal loss/neurodegeneration versus the marked cerebrovascular impact such as CAA. Aβ has a significant role in the pathogenesis of the two conditions, which frequently coexist with patient-specific burden of each pathology ^106^. One of the strongest phenotypes observed in BAM2 depleted mice is the formation of CAA, and it is possible that a differential functionality or survival of BAM2 versus microglia could play a role in the relative burden of CAA.

A recent publication reached the opposite conclusion regarding BAM protection against CAA ^107^. To generate CD36-deficient BAM cells, the authors used total body lethal irradiation and reconstitution with bone marrow from CD36 KO mice, an approach that generates concerns. For example, the presence of CD36 KO blood monocytes is expected in the reconstituted mice, obscuring the authors’ interpretation. Despite the technical issues, the age of the mice (one year old) used in the publication and our observations on older mice coincide in that NOX2-mediated ROS production is active in BAM from old AD mice.

Given the presence of BAMs in the perivascular space, choroid plexus, and meninges, but not in brain parenchyma, the Aβ buildup in the parenchyma in BAM2-depleted mice was counterintuitive. However, Aβ is thought to primarily be drained from the brain through large vessel perivascular spaces ^108–111^, the spaces where BAMs reside. The removal of the major phagocyte population from these clearance highways presumably sets off a cascade of events, whereby soluble Aβ oligomers build up in the vasculature and seed the deposition of overt CAA, creating a retrograde chain of events leading to the seeding of parenchymal plaques. This cascade is supported by our finding that both soluble and insoluble forms of Aβ were increased in BAM2 depleted mice brain-wide. BAMs have also recently been implicated in the flow dynamics of cerebral spinal fluid (CSF) through the glymphatic drainage system ^35^; thus, the buildup of plaque may also reflect changes in waste flow rate through the entire brain.

Microglial depletion using CSF1R inhibitors and a mouse strain with a targeted enhancer on the *Csf1r* gene ^44^ suggest that depletion of microglia may actually reduce amyloid buildup ^15–18,112^, with some studies showing a concomitant increase in CAA ^17,18^. However, it is clear that pharmacological CSF1R inhibition also depletes BAMs ^28,30,113^. With this in consideration, a unified idea as to why CSF1R inhibition causes decreased parenchymal plaques and increased CAA would be that BAMs and microglia are antagonistic regarding Aβ clearance: BAMs may be vital to amyloid clearance around blood vessels, and microglia may play a seeding role. When both are depleted, the parenchyma, dominated by microglia-dependent Aβ seeding, loses plaque, and the vasculature, dominated by BAM-dependent cleaning, gains plaque. However, when only BAMs are depleted, microglia Aβ seeding remains, and Aβ parenchymal buildup ensues without vascular clearance mechanisms.

Beyond an acceleration of disease in the absence of BAM2 cells, our work also highlights that BAM metabolic dysfunction may be a key contributor to AD progression. We demonstrate that BAM2 have a high rate of oxidative metabolism and produce high amounts of NADPH oxidase-generated ROS relative to microglia; this may be necessary for cells that reside in the waste clearance highway of the brain: the perivascular space. However, akin to T cells in the context of cancer ^114,115^, BAM2 in AD become hypofunctional, eventually reducing in number. Recent work highlights the importance of myeloid metabolic exhaustion in the context of AD and aging ^57,61,85,87,116^, the reversal of which may alone be therapeutic ^85,116^. While, as expected, we did detect metabolic changes in microglia during AD progression, our work suggests that BAM2 cells represent a subset of brain myeloid cells at the extreme regarding metabolic and general transcriptomic changes in AD. Single cell RNA Seq showed that microglia are a complex group of macrophages, but they appear to have in common a stability higher than the one that we observed in BAM ^117^.Our findings pave the way for future work focusing on therapeutic interventions to revive BAM mitochondrial flux, reduce excessive ROS, and even replace dysfunctional BAMs with new cells, as has recently been successfully implemented for microglia ^113,118,119^. Through preventing the breakdown of BAM2 function, we may in turn preserve the function of the neurovascular unit and limit the progression towards dementia.

---

**Figure S1.**
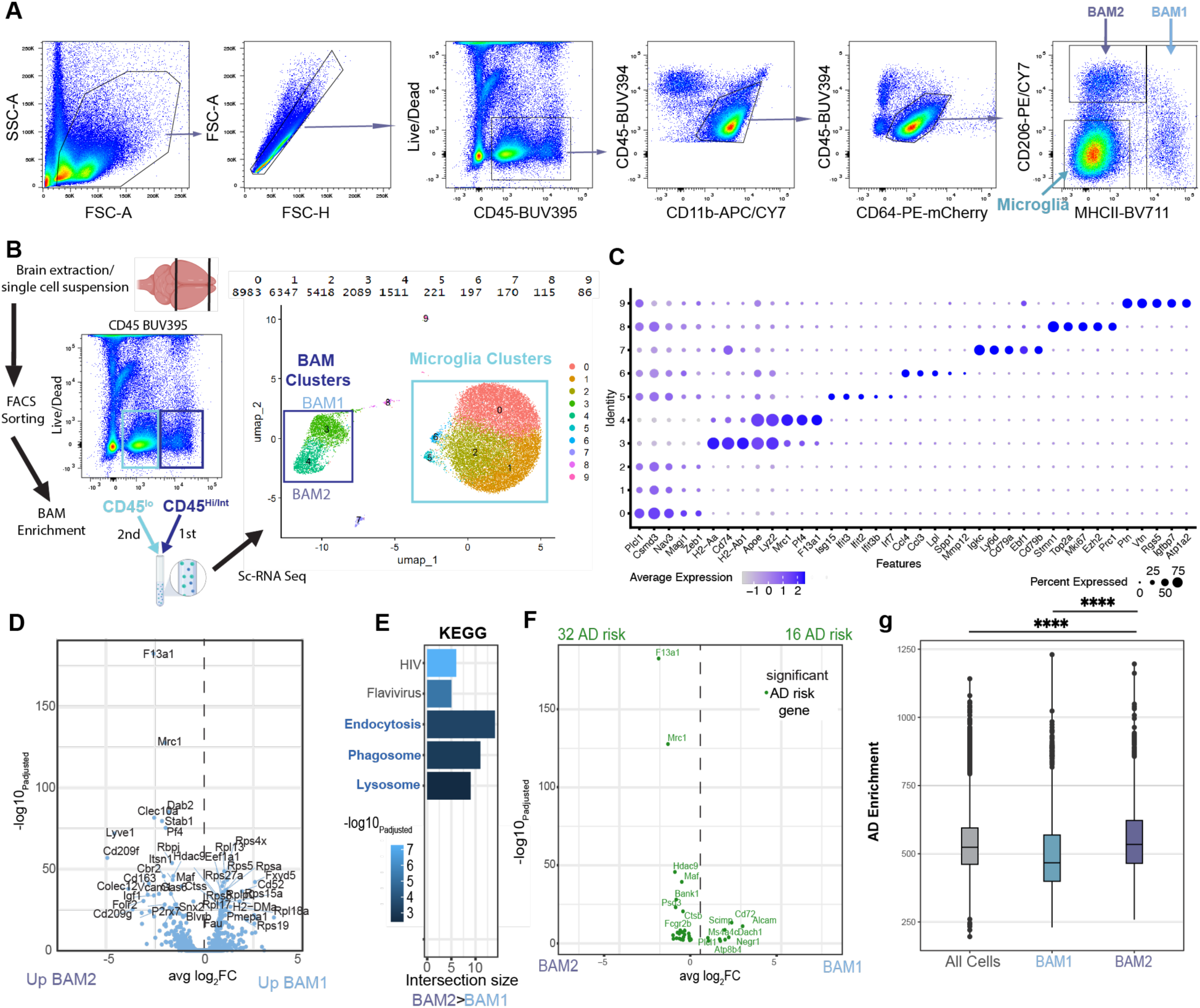
CD206^Hi^ BAMs (BAM2) are highly phagocytic and enriched in AD risk. Related to Figure 1. **A,** Gating strategy for MG and BAM subtypes. Brain resident macrophages were gated as CD45^+^CD11b^+^CD64^+^. Within this gate Microglia were gated as CD206^-^MHCII^-^, BAM2 were gated as CD206^+^MHCII^-^, and BAM1 were gated as MHCII+. **B**, Pipeline and BAM enrichment strategy for brain immune cell single cell RNA-Seq with corresponding UMAP defining cellular clusters and (above) number of cells within each cluster. **C**, Marker genes defining each cluster from (**B**). **D**, Volcano plot with all significant genes (P_adjusted_<.05) enriched in BAM2 vs BAM1. **E**, KEGG analysis of all pathways enriched in BAM2>BAM1. **F**, Whisker plot of all genes containing associated AD GWAS alleles (MONDO_0004975) enriched in BAM2 vs BAM1 clusters with (above) corresponding counts within each cell group. **G**, Per-cell AD enrichment score within BAM2, BAM1, and all cell aggregate clusters. Wilcoxon Rank Sum Test, P****<.0001. Inset box plots with median and 25^th^ and 75^th^ percentiles. UMAP= uniform manifold approximation and projection.

**Figure S2.**
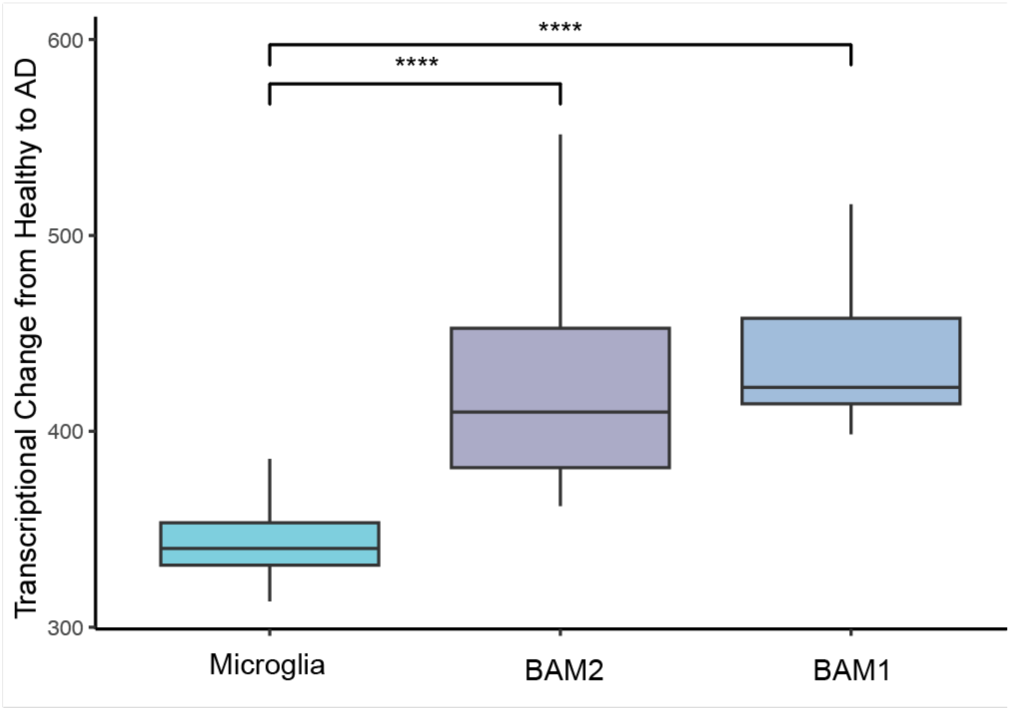
AD changes BAMs transcriptionally. Related to Figure 2. Quantification of the Euclidean distance of transcriptional change within brain macrophage subtypes between healthy and AD conditions with corresponding Chi-squared p-value. ****P<.0001, Pairwise t-test with Benjamini-Hochberg adjustment. Box plots= median + 25/75^th^ percentile.

**Figure S3.**
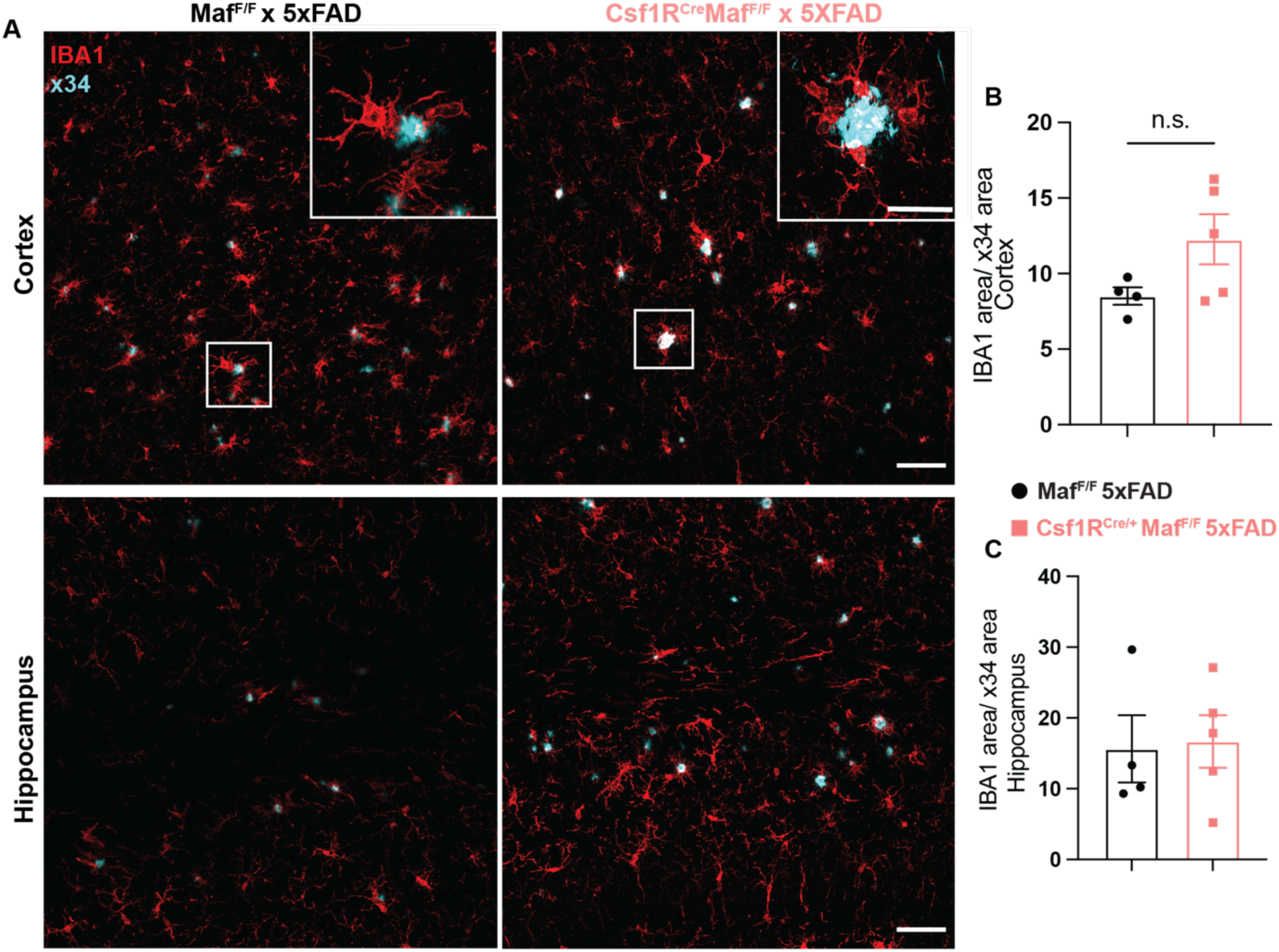
Microglia from Csf1R^Cre/+^*Maf*^F/F^ mice are reactive to amyloid. Related to Figure 3. **A,** IBA1^+^ (red) microglia staining and association with X-34 staining (cyan) of compacted amyloid in sections of BAM2-depleted Csf1R^Cre/+^*Maf*^F/F^ 5xFAD mice and *Maf*^F/F^ 5xFAD controls from similar regions of cortex and hippocampus. **B-C**, quantification IBA1 area normalized to amyloid area in (**B**) cortex and (**C**) hippocampus (n=4 *Maf*^F/F^ 5xFAD, 5 Csf1R^Cre/+^*Maf*^F/F^ 5xFAD mice). Two-tailed unpaired student’s t-test. Scale bar=100µm low magnification images, 50µm for high magnification insets.

**Figure S4.**
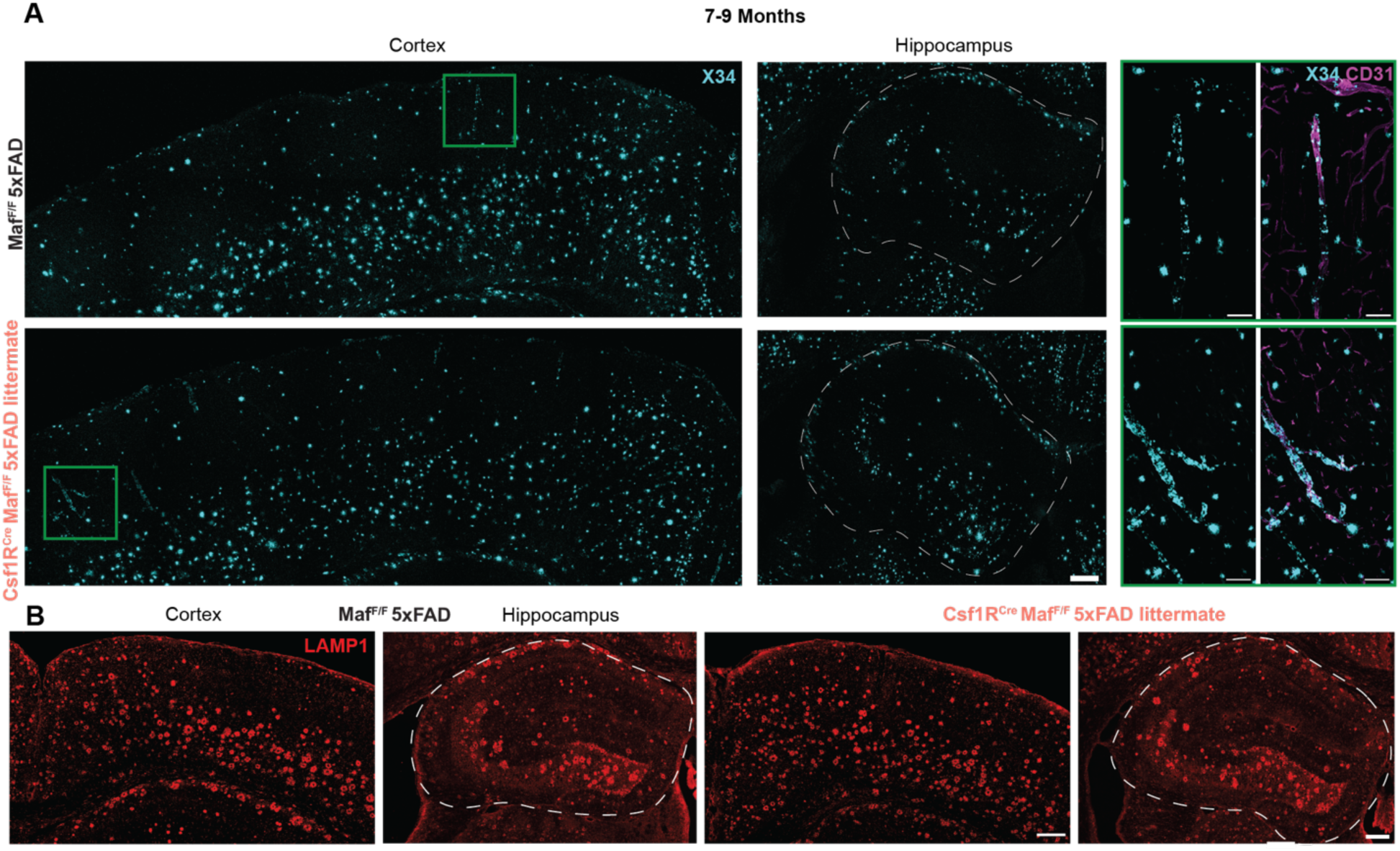
BAM2 depletion has minimal effect on Aβ load and neurodystrophy at late disease stages. Related to Figure 4. **A**. Representative micrographs of X-34 amyloid staining in cortex and hippocampus in 7-9 month old Cre^+^ (BAM2 depleted) and Cre^-^ 5xFAD littermates with regions of interest (ROIs, green boxes) depicting CAA in pial and penetrating CD31^+^ blood vessels in cortex. **B.** As with **A** but for LAMP1 neurodystrophy marker. Scale bars=200µm large images, 50µm high mag ROIs.

**Figure S5.**
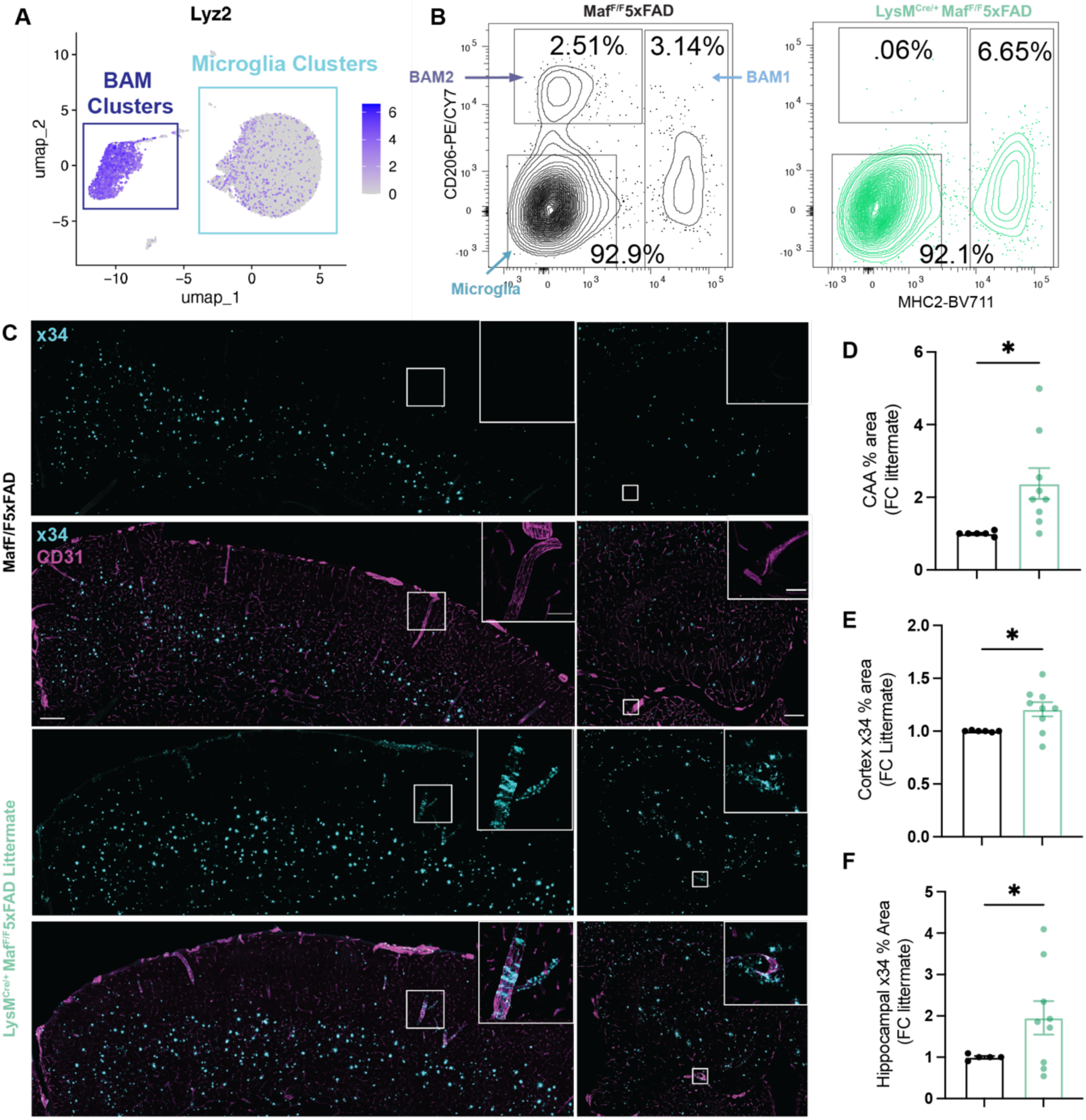
LysM^Cre/+^*Maf*^F/F^5xFAD have early acceleration of AD phenotypes. Related to Figures 3 and 4. **A,** Heatmap of relative *Lys2* (*LysM*) expression per cell in clusters detailed in **Fig. S1b**. **B**, Representative contour plots of brain macrophage populations (BAM2, top box; BAM1, right box; MG, bottom left box) with quantification of percentage of CD64^+^ gated cells. Note an increase in percentage of BAM1, which does not reflect a conversion of BAM2 into BAM1, but a rebound of BAM2, which are repopulated from monocytes ^32^. **C**, Representative micrographs depicting X-34 amyloid staining and CD31^+^ endothelial cells in LysM^Cre/+^*Maf*^F/F^ 5xFAD and *Maf*^F/F^ 5xFAD mice with high magnification insets depicting prominent CAA in LysM^Cre/+^*Maf*^F/F^ 5xFAD mice. **D-F**, Quantification of (**D**) CAA % blood vessel area, (**E**) cortical, and (**F**) hippocampal X-34 % area normalized to same sex littermate controls. Mean+/- S.E.M. Scale bars=200µm large images, 50µm insets.

**Figure S6.**
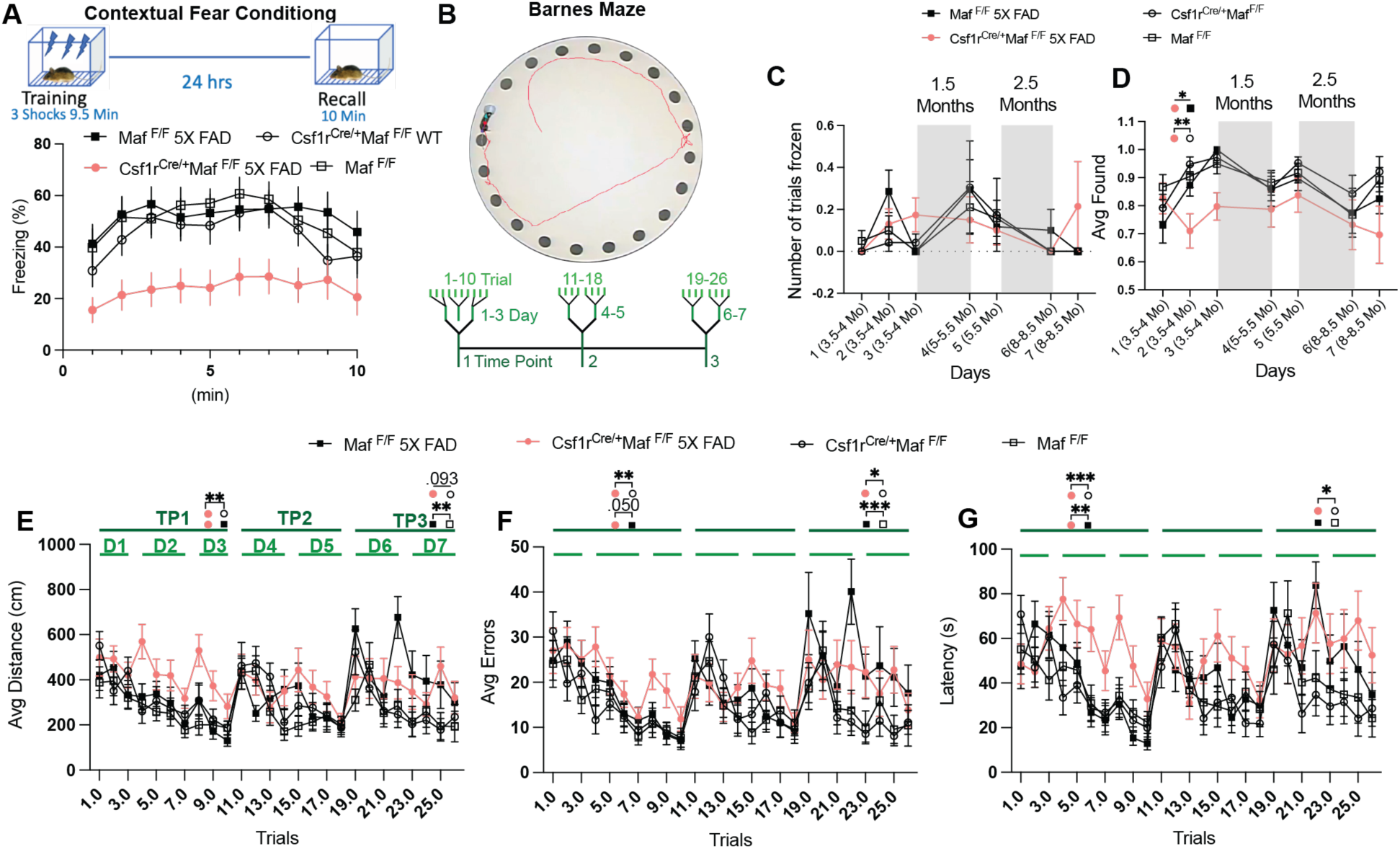
Behavioral analysis in BAM2 depleted AD mice. Related to Figure 4. **A**, Contextual Fear Conditioning experiment recall (**from** Fig. 4N) binned by 1-minute intervals. Significant 5xFAD x Minute interaction (Huynh-Feldt: F(6.982,432.880) = 2.397, p=0.021) and main effect of Cre (F(1,62) = 7.101, p=0.01). See main figure legend for n values. **B-G**, (**B**) Barnes Maze experiment (**from** Fig. 4P) with (**C**) average number of trials frozen per animal by day, (**D**) avg trials with hole found by day (Cre x 5xFAD x Day interaction (F(6,1858) = 3.766, p < 0.001) (**E**) avg Distance by trail, (**F**) avg Errors by trial, and (**G**) avg Latency per animal by trial. Two-way repeated measures ANOVA with Sidak’s multiple comparisons test. See *Materials and Methods* for n values. Significant pairwise comparisons and select p values within time points indicated by shapes associated with respective genotypes in the panel legend. Linear Mixed Effects Model revealed a main effect of Trial across Barnes Maze performance measures: (**D**) F(3,1856) = 17.854, p<0.001), (**E**) F(3,1856) = 13.585, p<0.001), (**F**) F(3,1856) = 9.188, p<0.001). (**C**) Sidak comparisons also revealed that Cre^+^5xFAD mice had a reduced ability to find the escape hole across days (all p<0.05), unlike all other groups. *P<.05,**P<.01, ***P<.001. TP=Time Point (months), D=days.

**Figure S7.**
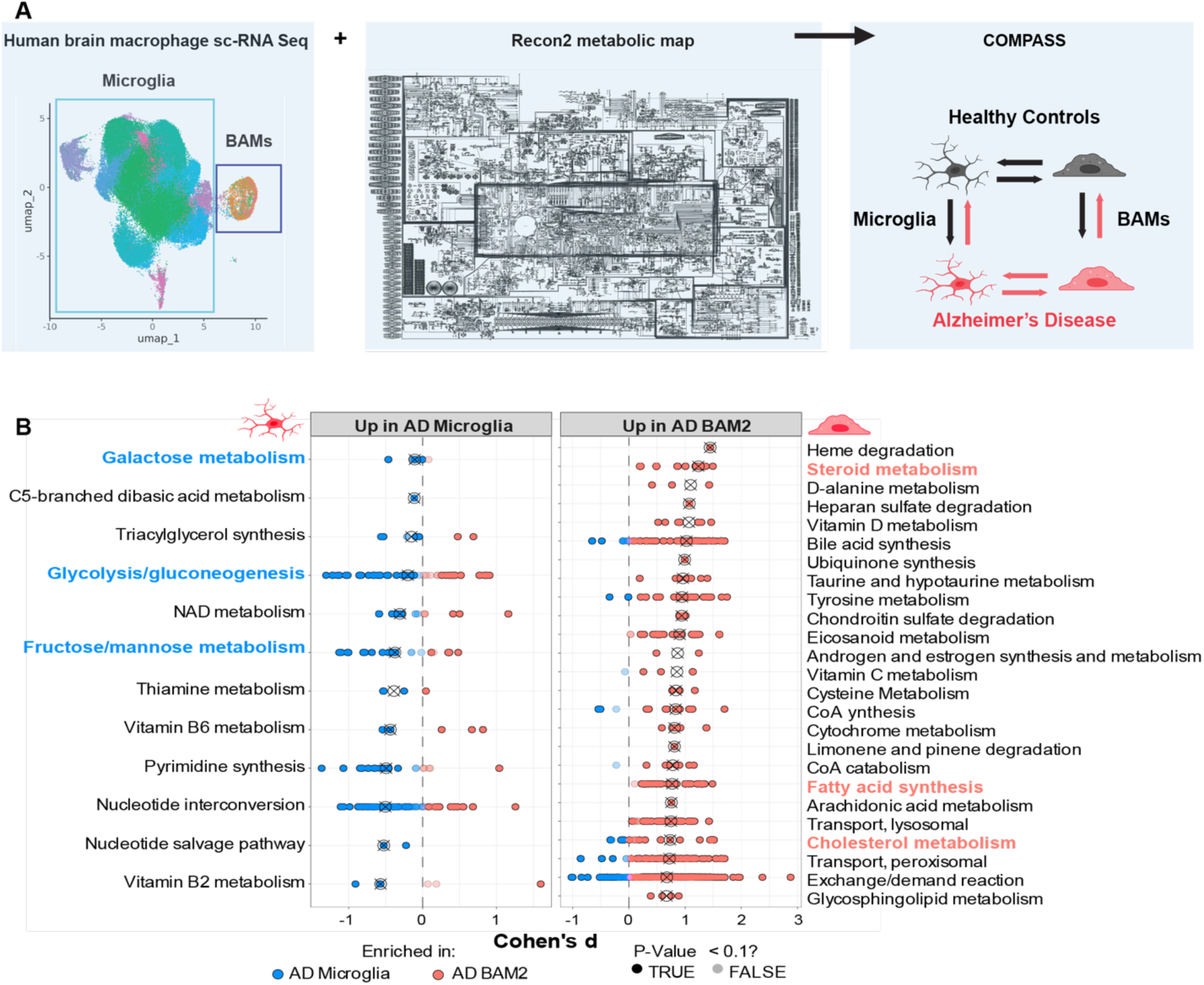
Human BAMs differ metabolically from microglia in health and AD. Related to Figure 6. **A,** Schematic of pipeline for *in silico* COMPASS analysis^54^ on aggregate human brain macrophages from control and AD patients. **B**, All discovered subsystems (Cohen’s d>.1) enriched in human microglia vs BAM2s from AD patients Colored subsystem names represent pathways discussed in main text. Circles with x=median enrichment for the respective subsystem. Individual circles=individual reactions within the subsystem that when shaded dark =P_adjusted_<.1 (Benjamini-Hochberg adjusted).

**Table S1. (separate file)**

Mouse Single-Cell RNA-seq comparisons.

**Table S2. (separate file)**

Human brain macrophage cluster markers.

**Table S3. (separate file)**

Compass individual reaction comparisons.

## Materials and Methods

### Mice

All animal protocols were reviewed and approved by the New York University Animal Care and Use Committee. C57BL/6J (000664), *LySM*^Cre^ (04781), and *Csf1r*^Cre^ (021024), 5xFAD (034840) strains were purchased from the Jackson Laboratories. *Maf*^FLOX^ strain was previously described^120^ and provided by C. Birchmeier. Male and female mice were used for all experiments. Ages used are specified in figures or legends. All litters were randomized into experimental groups.

### Drugs, chemicals, and recombinant proteins

70kDa Tetramethylrhodamine dextran (D1818, ThermoFisher) was dissolved in PBS and injected either retro-orbitally (.9 mg, 16 hours prior to perfusion) or into cisterna magna (5 µl, 10 mg/ml, 16 hours prior to perfusion). FAM-labeled beta-amyloid_1-42_ oligomers (AS-23526-01, Anaspec) were prepared by dissolving 250 µl ice cold 1,1,1,3,3,3-Hexafluoro-2-propanol (HFP, 105228, Sigma-Aldrich) for 1 hr in a chemical hood and transferring to a 1.5ml Eppendorf tube protected from light. After 1 hr the top of the tube was opened and evaporated overnight (ON). The next day Aβ1-42 HFP film was dissolved with 20 µl freshly opened anhydrous DMSO (570672) by scraping the sides of the tube with a pipette tip. Then 280 µl cold sterile PBS was added to the tube, vortexed for 10s, and incubated at 4C ON to form Aβ-derived diffusible ligands (ADDLs). The next day ADDLs were centrifuged at 14K G x 10 min at 4C and the supernatant was transferred to a new tube. The ADDL supernatant concentration was determined using BCA (Pierce) (23227, Thermo Fisher) and 1µM final concentration was added to single cell suspensions for 15 minutes at room temperature (RT). Methoxy X04 (4920, Tocris) was dissolved in DMSO and injected intraperitoneally (IP) into mice (10mg/kg)^58^ as a 1:5 DMSO:PBS solution. X-34 (74193, Cell Signaling) was dissolved in DMSO and 5µM was applied to slices (see IHC below). TMRE (564696, BD) was dissolved in DMSO and 40nM final concentration was used either alone or multiplexed with Aβ ADDLs for 15 minutes at RT. Carbonyl cyanide 40(trifluromethoxy)phenylhydrazone (FCCP, C2920, Sigma-Aldrich) was dissolved in DMSO and single cell suspensions were treated with 50 µM for 30 minutes followed by 15 minutes of Aβ ADDL/TMRE mixture. 500nM MitoSOX Red (M36007, Thermo Fisher) was prepared per manufactures instructions and added to pre-warmed single cell suspensions for 20 min at 37C. CellROX Deep Red (C10491, Thermo Fisher) was prepared per manufactures instructions and single cell suspensions were incubated at 500nM for 30 min at 37C prior to fixation. CellEvent Senescence Green (C10840, Thermo Fisher) was prepared per manufactures instructions and single cell suspensions were incubated 1:500 at 37C for 1 hour after fixation.

### Flow Cytometry

Mice were perfused with 5mM EDTA in PBS. All samples were kept on ice or at 4°C unless otherwise noted. Following perfusion, the olfactory bulbs, cerebellum, and brain stem were removed, and the remaining tissue was minced, and digested with collagenase D 240U/ml (11088858001, Sigma) in PFH solution: 2% fatty acid free BSA (700-107P-100, Gemini Bio), 1mM hepes buffer in PBS for 30 minutes at 37°C. Following digestion, a cell slurry was made with a 3mL syringe and tissue was mashed through a 100µM cell strainer. The single cell suspension was centrifuged at 1500 rpm for 5 minutes to pellet the cells. After centrifugation, the supernatant was aspirated, and the pellet was resuspended in 10ml 35% percoll in PFH (p4937, Sigma) and spun at 2000RPM for 30 minutes at RT with no break. After centrifugation, the supernatant containing myelin was aspirated, the cells were resuspended in cold PFH, filtered again, and centrifuged at 1500 rpm for 5 minutes. Cells were then incubated in CellROX Deep Red, TMRE, FCCP, DMSO, MitoSOX Red (concentrations and times above) or were immediately stained for following cell-surface markers with the following dyes/antibodies in PBS (if using Live/Dead e780 for live/dead) or PFH (if using DAPI for live/dead: Live/Dead e780 (1:1000, 65-0865-14, ThermoFisher), Rat anti-CD45 BUV395 or AF700 (1:200, 564729, BD, or 1:200, 103232, BioLegend), Rat anti-CD11b BV421 or APC/Cy7 (1:400, 101251, BioLegend or 1:400, 557657, BD), Mouse anti-CD64 BV650 or PE-Dazzle 594 (1:200, 740622, BD or 1:200, 139320, Biolegend), Rat anti-MHC2 BV711 (1:200, I-A/I-E, 107643 BioLegend), Rat anti-CD206 PE/CY7 (1:200, 141720, BioLegend) for 1hr at 4°C. Samples were then washed in PFH and DAPI was added before analyzing on a BD FACS Symphony A5. For samples stained with Cell Event Senescence Green (see above), samples were fixed after cell surface marker staining per manufacturer’s instructions.

Data was imported into FlowJo V10 (FlowJo LLC) for analysis. Microglia were gated Live/CD45^+^/CD11b^+^/CD64^+^/CD206^-^/MHC2^-^. CD206^High^ BAMs were gated as Live/CD45^+^/CD11b^+^/CD64^+^/CD206^+^/MHC2^-^. MHC2^+^ BAMs were gated as Live/CD45^+^/CD11b^+^/CD64^+^/^mid/low^/MHC2^+^ per^121^. All samples were compared to fluorescence minus one controls to determine % positivity and normalized MFI.

### Amyloid Beta ELISA

Mice were sacrificed with CO_2_ asphyxiation and brains were quickly removed; split in into hemispheres; the cerebellum, olfactory bulbs, and brain stem were removed; and the resulting brain was snap frozen in liquid nitrogen until they were ready to be used. To isolate the soluble brain fraction brains were homogenized in 0.2% diethanolamine in 50mM NaCl at a concentration of 100mg tissue/mL on ice using a sonicator. The resulting tissue was centrifuged at 100,000 x g for 1hr at 4°C. The supernatant (soluble fraction) was then neutralized by adding 1/10 volume of 0.5 M Tris HCL (pH 6.8) and vortexed gently and stored at -80°C. 200 µL of the pellet was mixed with 440 µL cold formic acid (minimum 95%, Sigma, 5-0507) in a microcentrifuge tube. Each sample was then sonicated for 1 minute on ice and 400ul was spun at 135,000 x g for 1 h at 4°C. 210 µL of the resulting supernatant (insoluble fraction) was diluted into 4 mL of room temperature fatty acid neutralization solution (1 M Tris base, 0.5 M Na 2HPO4, 0.05% NaN3, stored at room temperature) and mixed briefly and stored at -80°C. Soluble and insoluble fractions of each brain were thawed and run on the following ELISA kits for amyloid beta species (1-40 ThermoFisher KHB3481, 1-42 ThermoFisher KHB3441). Samples were normalized to their respective protein concentrations measured with a Bradford Assay (ThermoFisher 23200) and then to respective littermate controls.

### Mouse immunohistochemistry

#### Sample Preparation

Mice were perfused with 1x PBS followed by 4%PFA followed by ON post fixation in 4% PFA at 4C. After post fixation samples were immersed in 30% sucrose until they sunk and embedded in optimal cutting temperature compound (OCT) for slicing on a cryostat. 40 µm coronal slices were mounted on slides, skipping every other slice, from bregma ∼-1.22 to -2. Sections were rehydrated 2x 5 min in PBS followed by incubation for 2hrs in staining buffer: .1% triton x-100, .05% Tween-20, 2µg/mL heparin sodium salt (H3393, Sigma-Aldrich), .01% sodium azide in 1xPBS. Sections were incubated ON at 4C with the following primary antibodies Goat anti-CD31 (1:150, AF3628, R& D systems), Rabbit anti-IBA1 (1:1000, 019-19741, Wako), Guinea Pig anti-NeuN (1:1000, 266 004, Synaptic systems), Rat anti-CD206 (1:500, MCA2235, Bio-Rad), Rat anti LAMP1 (1:250, 1D4B, DSHB), Rabbit anti-Laminin (1:300, Thermo Fisher, PA116730). The next day samples were washed 3x 10 min with staining buffer and incubated for 2hrs at RT in the following secondary antibodies Donkey anti-Goat Alexa Fluor 488 (1:200, 705-545-003 JacksonImmuno), Donkey anti-Rabbit Alexa Fluor 568 (1:500, 711-575-152, JacksonImmuno), Donkey anti-Goat Alexa Fluor 647 (1:200, 705-607-003, JacksonImmuno) Donkey anti-Rat Alexa Fluor 647 (1:500, 712-605-152, JacksonImmuno), Donkey anti-Rat Alexa Fluor 568 (1:500, 712-575-153, JacksonImmuno) Goat anti-Guinea Pig Alexa Fluor 488 (1:500, A-11073, ThermoFisher). Samples were then washed 3x 10 min in staining buffer either mounted in Fluoromount G (00-4958-02, Thermo Fisher) or incubated for 10 min with 5µM X-34 (74193, Cell Signaling) in staining solution at RT followed by 3x 10 min washes and mounting.

#### Confocal microscopy

Images were acquired using a Zeiss LSM 800 or Zeiss 700 confocal microscope equipped with a 20x/.08 air Plan-Apochromat objective for X-34 % area quantification, LAMP1 % area quantification, CAA % area quantification, IBA1 area quantification, and NeuN Cell quantification. A 40x/1.3 N.A. oil emersion objective was used for Fig. 4A CAA images, Fig 4I LAMP1 images. A 63x plan apochromat N.A. 1.4 oil immersion Lense was used for Fig. S3 and S4 insets. Tiles were stitched using Zen Blue software (Zeiss). Identical microscope settings (e.g. imaging depth, z-step size, laser power, gain, optical zoom, offset, filters) were used for images in which fluorescent intensity or % area, were quantified and compared.

#### Image analysis

Images were imported into FIJI (ImageJ) for quantification. For X-34, IBA1, and LAMP1 % area quantification ROIs around motor and somatosensory cortex or hippocampus were drawn by hand. Images were z-projected (max intensity), background subtracted using the “subtract background” function (rolling ball radius 50 pixels), and filtered using the “median filter” function (pixel radius 2). Images were then thresholded identically for each fluorophore and the “analyze particles” function was used to calculate the total X-34, IBA1, or LAMP1 area per ROI which was divided by the total ROI area (*100) to calculate the % area for each ROI. For NeuN^+^ Cell counting, an experimenter blind to the experimental groups drew a single ROI around L5 for each sample and used the “cell counter” function to count individual NeuN^+^ neurons dividing the resulting number by the total area of the L5 ROI considered. For CAA quantification a single ROI was traced around all cortical pial and large penetrating CD31^+^ blood vessels (defined as blood vessels perpendicular to the pial surface) and CAA% area was calculated as the percent of CD31^+^ blood vessel area covered in X-34 using a custom Fiji macro: The CD31 channel was z projected (max intensity), background subtracted (rolling ball=50), median filtered (radius=3), thresholded (identically for each section compared), converted to a mask, eroded 2x with the “erode function”, and dilated 2x with the “dilate” function. The mask was selected with “create selection” which highlighted all CD31^+^ vessels and the total CD31 area was measured which represented the total vessel area considered. The X-34 channel was selected and the “restore selection” was applied. Within the restored section the X-34 area was thresholded (identically for compared samples), converted to a mask, and measured, which represented the total CAA area. The total CAA area was divided by the CD31 area (X 100) to generate the “CAA% area.” 2-6 slices were used per animal for each region for each analysis.

### Adipo-Clear

#### Sample preparation

The method was performed as described before^122^. Animals were anesthetized as described before, and an intracardiac perfusion/ fixation was performed with 1× PBS followed by 4% PFA. All harvested samples were postfixed in 4% PFA at 4°C overnight. Fixed samples were washed in PBS for 1 hour three times at RT.

#### Delipidation and permeabilization

Fixed samples were washed in 20, 40, 60, and 80% methanol in H2O/0.1% Triton X-100/0.3 M glycine (B1N buffer; pH 7) and 100% methanol for 1 hour each. Samples were then delipidated with 100% dichloromethane (DCM; Sigma-Aldrich) for 1 hour three times. After delipidation, samples were washed in 100% methanol for 1 hour twice and then in 80, 60, 40, and 20% methanol in B1N buffer for 1 hour each step. All procedures above were carried out at RT with shaking. Samples were then washed in B1N for 30 min twice followed by PBS/0.1% Triton X-100/0.05% Tween 20/heparin (2 µg/ml) (PTxwH buffer) for 1 hour twice before further staining procedures.

#### Immunolabeling

Half hemispheres of brain were incubated in Rabbit anti-Lyve1 (1:200, ab14917, abcam) and rat anti-CD206 (1:200, MCA2235, Bio-Rad) or Rabbit anti-IBA1 (1:300, 019-19741, Wako). Samples were incubated in primary antibody dilutions in PTxwH for 4 days at RT. After primary antibody incubation, samples were washed in PTxwH for 5 min, 10 min, 15 min, 30 min, 1 hour, 2 hours, 4 hours, and overnight and then incubated in secondary antibodies Donkey anti-Rabbit Alexa Fluor 568 (1:200,711-575-152, JacksonImmuno) or Donkey anti-Rat Alexa Fluor 647 (1:200, 712-605-152, JacksonImmuno) in PTxwH for 4 days at RT. Samples were lastly washed in PTwH for 5 min, 10 min, 15 min, 30 min, 1 hour, 2 hours, 4 hours, and overnight.

### Tissue Clearing

Samples were dehydrated in 25, 50, 75, 100, and 100% methanol/ H2O series for 1 hr for each step at RT. After dehydration, samples were washed with 50% MeOH/DCM followed by 2x 100% DCM for 1 hour each or until samples sunk to the bottom of their tube, followed by an overnight clearing step in dibenzyl ether (DBE; Sigma-Aldrich). Samples were stored at RT in the dark until imaging.

### Light Sheet Microscopy/ quantification

Hemispheres were mounted on a sample holder and submerged in an imaging chamber filled with DBE of a Zeiss light-sheet Z1 microscope equipped with a 5x N.A. 0.16 objective lens. 561 and 638nm lasers were used to capture Alexa 568 and 647 signal respectively. Lasers excited fluorophores sequentially on separate tracks and captured through LP585 and 660 filters respectively to minimize bleed through. Equal X/Y tiles and Z stacks were imaged from each hemisphere. To quantify the number of BAMs, 3D images were imported into fiji, max projected, background subtracted (rolling ball=50), median filtered (radius=2), thresholded equally, and the analyze particle function was used to identify BAMs in cortex. BAM number was normalized to the total cortical area considered per image. For microglia quantification, a single z slice from the 3D stack was used, and microglia were quantified as with BAMs.

### Mouse Single cell RNA seq

#### Sample preparation and sorting

Single cell suspensions were generated as for flow cytometry above from 4 5-6 month old WT mice (2M, 2F) with the following modifications. To prevent artifactual transcriptional signatures induced by cell processing, mice were perfused with 20 an inhibitor cocktail of 5µg/mL actinomycin D (A9789, Sigma) and 10µM triptolide (T3652, Sigma) in PBS with 5mM EDTA^123^. Following perfusion olfactory bulbs, cerebellum, and brain stem were removed and the remaining tissue was minced and digested with collagenase D 240U/ml (11088858001, Sigma) in PFH solution: 2% fatty acid free BSA (700-107P-100, Gemini Bio), 1mM hepes buffer in PBS, with 5ug/mL actinomycin D, 10µM triptolide, and anisomycin 27.1 µg/ml (A9789, Sigma) for 30 minutes at 37°C. Cells were stained with Rat anti-CD45 BUV395 and washed as above. Individual animals were labeled using separate Cell Multiplexing Oligo labels (Protocol 4, CG000391, 10x Genomics), prior to pooling samples and incubated with DAPI. The single pooled sample from all animals was sorted into CD45^Hi^ and CD45^Lo^ populations (See Fig. 1c) using a FACSAria II Sorter (BD).

### Library construction and sequencing

scRNA-seq and Cell Surface Protein libraries were constructed using the Chromium Single Cell 3′ v3.1 Reagent Kit (10x Genomics, PN-1000268) according to the manufacturer’s protocol. Briefly, cell populations were stained with the 3ʹ CellPlex Kit Set A (10x Genomics PN-1000261) and counted using a BioRad TC20 Automated Cell Counter. Approximately 19K CD45^Hi^ cells and 40K CD45^Lo^ cells were pooled and placed in a channel of a Next GEM Chip G (10x Genomics, PN-1000120), loaded onto the Chromium Controller (10x Genomics), and single cells were encapsulated into emulsion droplets. Reverse transcription and library preparation were performed on a C1000 Touch Thermal cycler with 96-Deep Well Reaction Module (Bio-Rad). cDNA was amplified 11 cycles and evaluated on a Agilent BioAnalyzer 2100 using a High Sensitivity DNA Kit (Agilent Technologies). Final Gene Expression and Cell Surface Protein libraries were amplified 14 and 6 cycles respectively and visualized on an Agilent TapeStation 4200 using High Sensitivity D1000 ScreenTape (Agilent Technologies). Individual libraries were diluted to 2nM and pooled for sequencing. Pools were sequenced across 10% of an S4 200 cycle flowcell (28bp Read1, 10bp Index1, 10bp Index2 and 91bp Read2) on the NovaSeq 6000 Sequencing System (Illumina).

### Data processing and analysis up to UMAP

Raw sequencing data, including gene expression, unique molecular identifiers (UMIs), and multiplexing oligos, were processed using 10X Genomics CellRanger^124^. The processed data was further analyzed using R version 4.3.1. We excluded any cells with ≤ 300 RNA counts or ≥ 5% mitochondrial reads. Cell doublets and negatives were identified and excluded from downstream analysis using the HTODemux function^125^. Transcriptomic data was normalized using a log normalization transformation. We clustered the cells based on the top 2,000 variable genes from their transcriptomic profiles and visualized this clustering using UMAP projections of the transcriptomic features^126^.

### AD Enrichment Scores

As a proof of principle, we assessed whether certain cell populations were enriched for genes relevant to Alzheimer’s Disease. We performed single-cell gene set enrichment analysis (GSEA) for each of the main clusters and subclusters using genes from the NHGRI-EBI Alzheimer’s Disease GWAS gene set (MONDO_0004975). The enrichment scores were generated using the escape tool^127^, a package designed for single-cell GSEA.

### Human AD postmortem samples

#### Sample Details

| Diagnosis | Case | Age (Yrs) | Sex | ABC Score |
| --- | --- | --- | --- | --- |
| AD | TN12-78 A | 67 | M | A3, B3, C3 |
| AD | TN11-54 A | 68 | U | A3, B3, C3 |
| AD | TN16-22 A | 86 | F | A3, B3, C3 |
| AD | TN15-86 A | 81 | U | A3, B3, C3 |
| AD | TN16-43 A | 59 | U | A3, B3, C3 |
| AD | TN16-6 A | 73 | F | A3, B3, C3 |
| AD | TN14-23 A | 55 | F | A3, B3, C3 |
| AD | TN12-73 A | 93 | F | A3, B3, C3 |
| AD | TN13-2 A | 91 | F | A3, B3, C3 |
| AD | TN12-32 A | 70 | U | A3, B3, C3 |
| AD | TN16-19 A | 85 | F | A3, B3, C3 |
| AD | TN13-64 A | 68 | F | A3, B3, C3 |
| AD | TN17-52 | 72 | F | A3,B3,C3 |
| AD | TN17-27 | 104 | F | A3,B3,C3 |
| AD | TN17-18 | 73 | F | A3,B3,C3 |
| AD | TN16-122 | 73 | M | A3,B3,C3 |
| Control | TN18-10 | 76 | M | A1,B1,C0 |
| Control | TN17-127 | 98 | F | A0,B2,C0 |
| Control | TN18-21 | 91 | M | A1,B2,C0 |
| Control | TN16-79 | 97 | F | A0,B2,C0 |
| Control | TW18-27 | 96 | F | A0,B1,C0 |
| Control | TN15-56 | 45 | F | FOCAL CORTICAL DYSPLASIA, TYPE IA |

#### Sample preparation/CODEX

Tissues were fixed in 4% PFA at 4°C for 24 hours, dehydrated through graded ethanols and xylene, then infiltrated with paraffin (Paraplast X-tra, Leica cat 39603002) on a Leica Peloris II tissue processor and embedded on a Leica Arcadia embedder.

7 µm sections were immunostained on a Leica BondRx auto-stainer according to the manufacturer’s instructions. In brief, sections were deparaffiinized online and treated with 3% H_2_O_2_ to inhibit endogenous peroxidases, followed by antigen retrieval with either ER1 (Leica, AR9961; pH6) or ER2 (Leica, AR9640; pH9) retrieval buffer at 100o for either 20 or 60 minutes. After blocking with Primary Antibody Diluent (Leica, AR93520), slides were incubated with the first primary antibody and secondary HRP polymer pair, followed by HRP-mediated tyramide signal amplification with a specific Opal® fluorophore. Once the Opal® fluorophore was covalently linked to the antigen, primary and secondary antibodies were removed with a heat retrieval step. This sequence was repeated 5 more times with subsequent primary and secondary antibody pairs, using a different Opal fluorophore with each primary antibody (see table below for primary antibody sequence and reagent details). After antibody staining, sections were counterstained with spectral DAPI (Akoya Biosciences, FP1490) and mounted with ProLong Gold Antifade (ThermoFisher Scientific, P36935). Experimental details for two panels are below:

*Panel 1*

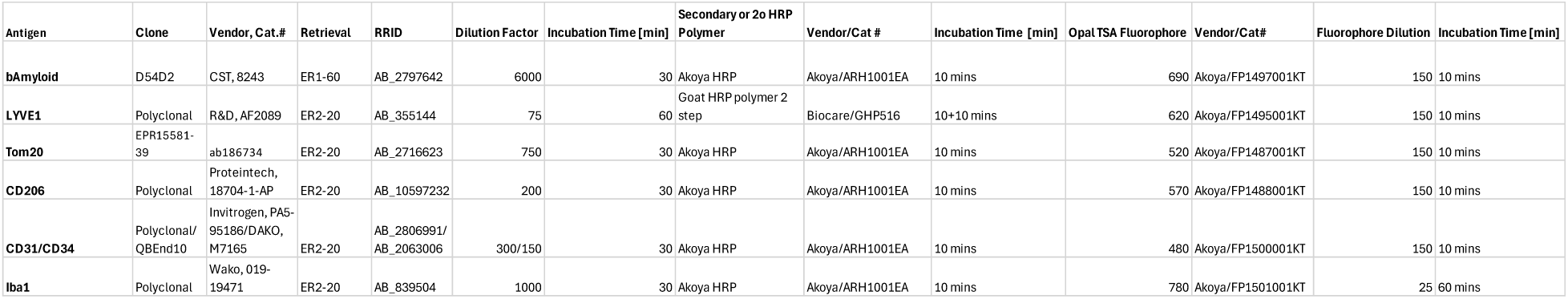

*Panel 2*

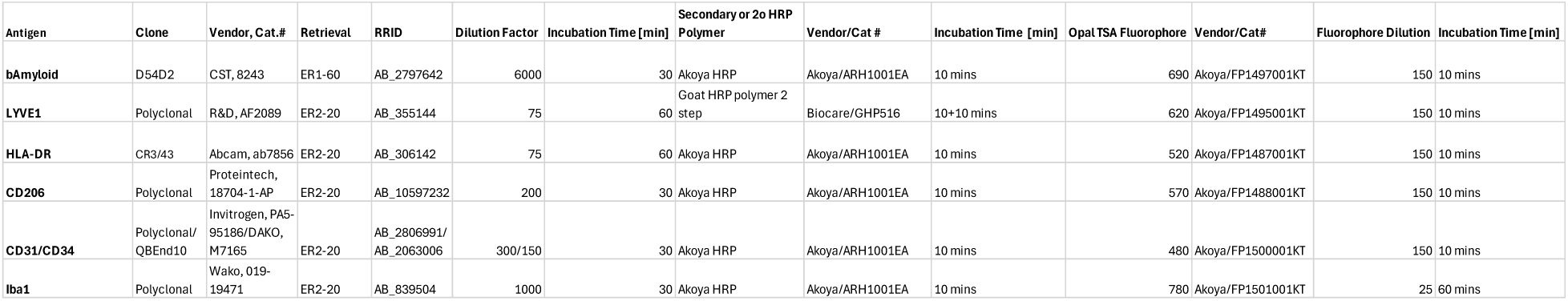

Semi-automated image acquisition was performed on an Akoya Vectra Polaris (PhenoImagerHT) multispectral imaging system. Slides were scanned at 20X magnification using PhenoImagerHT 2.0 software in conjunction with Phenochart 2.0 and InForm 3.0 to generate unmixed whole slide qptiff scans. Image files were uploaded to the NYUGSoM’s OMERO Plus image data management system (Glencoe Software).

### Human AD in silico analyses

#### Preprocessing and integration of human scRNAseq datasets

We leveraged two previously published single-nucleus RNAseq datasets of human AD patients and healthy controls from the ROSMAP project for our analysis. From the multiregion dissection data, we exclusively filtered the single-cell matrix to cells annotated by the original authors as CNS-associated macrophages. From the DLPFC-2 data, we separately filtered out both macrophages and microglia. We merged these three single-cell structures together, annotated them with the metadata provided by ROSMAP’s clinical manifest, and integrated them using Seurat’s CCA integration approach and regressing out the dataset of origin during SCTransform.

### Annotation and cluster enrichment analysis

Single-cell data was clustered using the standard Seurat workflow after SCTransform to generate several microglial clusters and one macrophage supercluster. Macrophages were further subclustered by subsetting and rerunning SCTransform on the supercluster, again regressing out dataset of origin. CD206-hi and MHCII-hi macrophage clusters were identified by computing enrichment of CD206+ and MHCII+ signatures previously determined from analysis of single-cell murine data. We exponentiated the signature scores produced by Seurat’s AddModuleScore utility and took the ratio of the CD206+ score to the MHCII+ score to identify whether a given cluster was more strongly defined by the gene program of one state or the other. Subclustering also identified a minor population of dendritic cells, defined by expression of Itgax and Zbtb46.

To compute the relative enrichment of CD206-hi and MHCII-hi macrophages in AD and healthy donors, we calculated the total number of macrophages from all human donors in each cluster, partitioned by AD status, and defined a naïve enrichment score as the fraction of macrophages in a given cluster originating from AD patients divided by the total size of the cluster. To normalize for overall differences in macrophage numbers, we normalized these enrichment scores by dividing by the mean enrichment score across all clusters.

### Metabolic flux modeling

We computed metabolic flux through all reactions annotated in RECON2 using Compass, a flux-balanced analysis algorithm that uses mixed integer programming to evaluate the utilization of metabolic reactions, constrained by stoichiometry and weighted by RNA expression of enzymes. We randomly pseudobulked the single-cell counts matrix for macrophages alone or for both macrophages and microglia, aggregating 10 cells into each metacell, to improve estimate robustness. We provided these matrices as inputs for Compass against the human reference. The resulting penalties were converted to flux estimates by taking the negative logarithm of one plus the penalty, and fluxes were then min-fixed and Z-scored. The effect size for the change in flux for each reaction was calculated as the Cohen’s d, or the mean divided by the standard deviation. These effect sizes were used for downstream visualization.

### Computation of transition distance from healthy to AD samples

To create a quantitative measure of the extent to which cell types change their transcriptional and metabolic state during AD pathogenesis and progression, we calculated the Euclidean distance between the transcriptomes and Compass penalty estimates in microglia, CD206-hi BAMs, and MHCII-hi BAMs between healthy and AD donors. We compute this pairwise distance between every single pair of cells in healthy and AD donors, as samples are not formally paired. To make this computation more efficient, we calculated the square of the distance 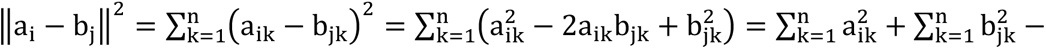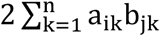, where a_i_ and b_j_ are the counts or penalties for one cell or pseudobulked cell in AD and healthy patients, respectively. We then take the square root of this value to arrive at the final Euclidean distance. We then plot the distribution of these distances in each cell type as a boxplot to determine which cell types show the strongest transitions during disease progression.

### Linear models of macrophage frequency in human donors

We merged ROSMAP clinical data with our annotated single cell dataset to generate a unified structure with both cell annotation and patient of origin metadata. We computed the fraction of cells per patient that were annotated as BAMs and defined a CD206:MHCII enrichment score, 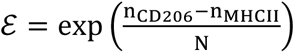, in which n_MHCII_ is the number of CD206-hi BAMs, is the number of MHCII-hi BAMs, and N is the total number of cells. We filtered to only samples in which at least 0.1% of cells were annotated as macrophage in order to remove patients in which cell capture was insufficiently well distributed across clusters. We then built linear models in which either BAM fraction or CD206:MHCII enrichment score are predicted by a linear combination of sex, Braak score, postmortem interval, CERAD score, educational attainment, E2 allele count, and E4 allele count. We plotted the coefficients and p-values associated with each of these predictors as a forest plot.

### Sources of data for human in silico analyses

1. DLPFC-2 (465 donors): Project SynID syn2580853.
2. Ref ^64^
3. Multiregion (48 donors, 6 regions, 283 samples) ^63^

### Behavioral Procedures and Analysis

#### Contextual Fear Conditioning

Mice are subjected to a two-trial protocol wherein they receive brief unconditioned stimuli (foot-shocks) paired with a novel context (conditioning chamber) - contextual fear memory is assessed by quantifying the percentage of freezing behavior upon re-exposure to the same context. On day 1 (“training day”) mice were brought in their home cages to the fear condition room of the NYUMC Rodent Behavior Laboratory and allowed to acclimate for 1 hour. Following acclimation, mice were placed in individual fear conditioning chambers (Habitest, Coulbourn Instruments) and allowed to acclimate for 180s. Mice were then shocked three times using 0.3mA of current for 2s duration, with each shock initiated at 180s, 330s, and 540s. Mice where then removed from the chambers at 570s and placed immediately back in their home cage.

Approximately 24 hours later (day 2, “testing day”) mice were brought back to the fear conditioning room and allowed to acclimate for 1 hour. Mice were then placed in the same conditioning chamber as on day 1 for 5 minutes. The amount of time frozen was automatically quantified for each animal using FreezeFrame (version 3) software, with freezing bout threshold set to 8 (motion/pixel noise) and 1.75s (minimum freeze duration). Chambers were cleaned with 70% ethanol and thoroughly dried prior to testing each mouse on both training and testing days.

### Barnes Maze

The Barnes maze assesses visuospatial learning and memory in mice. Mice were tested based on our protocols as previously published in the NYUMC Rodent Behavior Laboratory (Eugenio Gutiérrez-Jiménez et al., 2024; Canepa et al., 2023; Lin et al., 2020). The Barnes maze apparatus comprised a circular beige platform 91.5 cm in diameter positioned 91.5 cm above the floor, with 20 holes (each 5 cm diameter) evenly spaced near its edge (2.5 cm from perimeter) (San Diego Instruments). An escape box (10 × 8.5 × 4 cm) was placed under a designated hole (target escape hole) for each mouse at each age timepoint. Animals were motivated to find the escape hole to avoid the open space, bright lighting and white noise generator (600 lux and 72 dB at platform surface, respectfully) mounted above the maze.

On each testing day, mice were brought to the testing room to acclimate for 1 hour prior to testing. On the first day of testing at 3.5-4 months of age mice were given an initial habituation trial which involved guiding them to the target hole from the center of the table over the course of 30 seconds using an inverted transparent glass beaker. On subsequent standard training trials, each mouse was placed in the center of the maze within an opaque acrylic start box (15 × 15 × 20 cm), with the white noise stimulus applied. The box was lifted, and the animal was given a 2-min window to locate and enter the designated escape hole. Upon successful entry, the white noise was turned off and the mouse returned to the home cage. If the mouse failed to find the escape hole within the allotted time, it was gently guided to the target hole beneath the inverted beaker, as per habituation trials. The platform and escape box were cleaned with 70% ethanol and dried between each mouse and the maze platform was rotated three holes clockwise after all mice completed a trial. Two-minute probe trials were conducted, identical to standard trials, except that the escape box was removed.

In total, mice completed up to 26 trials (10 trials across 3 days at time point 1 (3.5-4 months), and 8 trials across 2 days at time points 2 (5-5.5 months) and 3 (7.5-8 months) (**see Fig. 4N**). The designated escape hole was varied at each age time point. Trials in which mice froze in the center of the arena and failed to search for more than 40s were repeated up to three times after which the entire trial was eliminated. These circumstances represented only 3% of total trials and there was no clear trend between genotype (**see Fig. S6C**). All mice completed the first time point – a small proportion of animals died during testing or were used for other purposes before finishing the last two time points. Different mice were used for Contextual Fear Conditioning and Barnes Maze testing. N values per genotype and time were as follows:

Time point 1: n=10 WT males, 10 WT females, 11 Cre^+^ males, 13 Cre^+^ females, 8 5xFAD males, 13 5xFAD females, 12 Cre^+^ 5xFAD males, 11 Cre^+^ 5xFAD females

Time point 2: n=10 WT males, 9 WT females, 9 Cre^+^ males, 12 Cre^+^ females, 7 5xFAD males, 10 5xFAD females, 9 Cre^+^ 5xFAD males, 11 Cre^+^ 5xFAD females

Time point 3: n=6 WT males, 8 WT females, 9 Cre^+^ males, 10 Cre^+^ females, 4 5xFAD males, 6 5xFAD females, 6 Cre^+^ 5xFAD males, 8 Cre^+^ 5xFAD females

Each trial was recorded using an overhead camera, and videos were analyzed with EthoVision XT tracking software (version 17, Noldus Information Technology, Inc.). Outcome measures included whether they located the escape box (found [1/0]), the time to locate it (latency [s]), the number of non-target holes visited (errors [#]) and the distance traveled before entering the hole (distance [cm]). Data from the Barnes maze were analyzed using a generalized linear mixed-effects model (random intercept per mouse with variance components covariance structure and repeated measures factors [Age/timepoint, Day and Trial] modeled with an autoregressive (AR1) residual covariance structure) to test the effects of Sex, Cre status, AD genotype, and testing Age and their interactions, as well as within-session learning across Days and Trials. Latency, Distance and Errors were modeled using a normal distribution and identity link function. Found was modeled using a binary distribution with a logit link function. Pairwise comparisons and estimated marginal means were calculated using the Sidak-adjusted significance level (α = .05) to test significant main effects and interactions.

## Statistical Analysis

Statistical analysis and data visualization were performed in GraphPad Prism (v10), R studio, or SPSS. Details regarding tests used and number of replicates for each experiment are described in figure legends or respective methods section. Statistical significance was determined by two-tailed unpaired Student’s t-test when comparing two independent groups or paired t-test when comparing values from the same group. Wilcoxen signed-rank test was used in paired sequencing data when normality could not be assumed, grouped data were analyzed by two-way ANOVA or linear mixed-effects model. Individual group differences were determined by Tukey’s post hoc test for two-way ANOVAs or Sidak’s post hoc test for repeated measures mixed-effects model with α set to .05. Significant interactions are reported in figure legends. Outliers were removed using the ROUT method with Q Set to 0.1%. All data are presented as individual values and mean ± S.E.M or boxplots with maximum values, quartile ranges, and median value.

## Code Availability

Any additional information required to reanalyze the data reported in this paper is available from the lead contact upon request.

## Acknowledgments

We thank Michael Pacold, Daniel Mucida and Shohei Tavazoie for many helpful discussions. We thank Michael Cammer from NYU Microscopy Core. The Microscopy Core is partially supported by NYU Cancer Center Support Grant NIH/NCI P30CA016087 at the Laura and Isaac Perlmutter Cancer Center, S10 RR023704-01A1 and NIH S10 ODO019974-01A1. We thank the Cytometry and Cell Sorting Laboratory (Peter Lopez and Catherine Rapelje). We thank Cindy Loomis from NYU Experimental Pathology Core for expertise and assistance with CODEX and the quality control of each antibody. We thank Thomas Wisniewski and Arlene Faustin for the human autopsy samples. EK was supported by R01 NS122316. DA was supported by NIH 5T32MH019524-31 and NIH 5T32NS086750-08. Funded by NIH R01AG068142 (JJL). Early work on myeloid cells in the brain (during EAE) was supported by National Multiple Sclerosis RG 4299-A-5 (JJL).

## Author Contributions

Conceptualization: DA, JHL, HMS, JJL

Methodology: DA, NPR, ACM, ZZG, EGL, YM, JHL, HMS

Investigation: DA, NPR, ACM, ZZG, EC, YM, JHL

Visualization: DA, AM, ZZG,

Funding acquisition: DA, JJL, EK Supervision: DA, WBG, EK, HMS, JJL Writing – original draft: DA

Writing – review & editing: DA, AM, ZZG, BGL, YM, WBG, ACM, JHL, HMS, JJL

